# Opposing roles for TGFβ- and BMP-signaling during nascent alveolar differentiation in the developing human lung

**DOI:** 10.1101/2023.05.05.539573

**Authors:** Tristan Frum, Peggy P. Hsu, Renee F.C. Hein, Ansley S. Conchola, Charles J. Zhang, Olivia R. Utter, Abhinav Anand, Yi Zhang, Sydney G. Clark, Ian Glass, Jonathan Z. Sexton, Jason R. Spence

## Abstract

Alveolar type 2 (AT2) cells function as stem cells in the adult lung and aid in repair after injury. The current study aimed to understand the signaling events that control differentiation of this therapeutically relevant cell type during human development. Using lung explant and organoid models, we identified opposing effects of TGFβ- and BMP-signaling, where inhibition of TGFβ- and activation of BMP-signaling in the context of high WNT- and FGF-signaling efficiently differentiated early lung progenitors into AT2-like cells *in vitro*. AT2-like cells differentiated in this manner exhibit surfactant processing and secretion capabilities, and long-term commitment to a mature AT2 phenotype when expanded in media optimized for primary AT2 culture. Comparing AT2-like cells differentiated with TGFβ-inhibition and BMP-activation to alternative differentiation approaches revealed improved specificity to the AT2 lineage and reduced off-target cell types. These findings reveal opposing roles for TGFβ- and BMP-signaling in AT2 differentiation and provide a new strategy to generate a therapeutically relevant cell type *in vitro*.

## Introduction

During lung development, all epithelial cells of the pulmonary airways and alveoli differentiate from specialized progenitor cells that reside at the tips of a tree-like network of epithelial tubes, called bud tip progenitors (BTPs)^1–3^. During the pseudoglandular stage of development, BTPs undergo repeated bifurcations, a process known as branching morphogenesis, giving rise to the conducting airways (bronchi, bronchioles). Later, during the canalicular stage, cells of the alveolar epithelium begin to differentiate. Cells located within the epithelial stalk region directly adjacent to BTPs begin to express alveolar type 1 (AT1) marker genes, while BTPs begin to express markers consistent with alveolar type 2 (AT2) differentiation^2–6^. How descendants of BTPs are influenced to differentiate into airway or alveolar cell fates is determined by cues from their environment, but the mechanisms promoting human alveolar differentiation are not fully characterized^7–9^.

Clinical data has shown that treating premature infants with dexamethasone (a glucocorticoid receptor (GR) stimulating hormone) and/or inducers of cAMP-signaling promotes lung epithelial maturation and surfactant production^10–12^. This information has been leveraged to develop methods to differentiate primary or iPSC derived lung epithelium into AT2-like cells^13–17^. Interestingly, studies in mice have shown that GR-signaling is not required for alveolar cell fate specification, but rather loss of GR leads to a reduced size of the alveolar compartment^18, 19^. Likewise, AT2 differentiation is only moderately reduced in mice lacking the primary effector of cAMP-signaling^20^. These results suggest that while GR- and cAMP-signaling can play an important role in alveolar cell maturation and surfactant production, other cell signaling pathways likely operate alongside GR and cAMP-signaling to promote AT2 differentiation from BTPs.

More recently, single cell characterization of the developing human lung has been applied to identify factors that regulate human BTPs and their differentiation^5, 6, 21–23^. Here, we focus on cell signaling events that occur during nascent alveolar differentiation in the developing human lung. We leveraged single cell RNA sequencing (scRNA-seq) data from human fetal lungs and used computational approaches to interrogate the signaling events that take place between BTPs and RSPO2+ mesenchymal cells, which comprises a major component of the BTP niche^21^. We also developed and interrogated a serum and growth factor-free human fetal lung explant system that undergoes nascent alveolar differentiation. Collectively, data from these analyses point to TGFβ- and BMP-signaling as important cell signaling pathways that work in opposition to promote AT2 differentiation, with low levels of TGFβ- and high levels of BMP-signaling associated with differentiation of BTPs to AT2 cells.

We tested this model in BTP organoids^3^, combining simultaneous TGFβ-inhibition and BMP-activation in the presence of activated WNT- and FGF-signaling to efficiently induce AT2 differentiation in BTP organoids. AT2-like organoids generated using this method maintain an AT2 phenotype when expanded in serum-free monoculture conditions optimized for primary adult human AT2 organoids. As a comparison, we also applied a Dexamethasone and cyclic AMP (CK + DCI) protocol used commonly for generating iPSC-derived AT2-like cells^17, 24, 25^, and found that CK + DCI treated BTP organoids gave rise to AT2-like cells, albeit with reduced specificity. Nonetheless, CK + DCI induced AT2 cells could also be expanded in primary AT2 organoid media, facilitating three-way comparison between AT2-like organoids produced by both methods and benchmarked against primary adult AT2 organoids in the same media. This analysis revealed that BTP organoids differentiated with modulation of TGFβ- and BMP-signaling maintain a more homogenous population of AT2-like cells than BTP organoids differentiated with CK + DCI. Of note, comparison of AT2-phenotype retaining cells induced by both methods revealed highly similar AT2-like cells based on scRNA-seq data.

Taken together our findings identify TGFβ- and BMP-signaling as important pathways regulating nascent alveolar differentiation *in vivo* that can be leveraged to generate AT2-like organoids, which produce and secrete lamellar bodies, and have long-term expansion capabilities when grown in media optimized for primary AT2 organoids.

## Results

### High BMP- and low TGFβ-signaling are associated with alveolar type 2 cell differentiation during lung development

Using our previously published scRNA-seq data from human fetal lungs^23^ we applied CellChat^26^ to computationally interrogate cell-cell communication between BTPs and RSPO2-positive mesenchyme, which surrounds BTPs and is an established source of BTP niche cues in humans^21^. To identify interactions enriched in BTPs, we also included non-BTP distal epithelial cells (also referred to as ‘bud tip adjacent’, or stalk cells) and their associated mesenchyme population, identified by co-expression of SM22, ACTA2 and NOTUM^21, 27, 28^. This analysis predicted that RSPO2-positive mesenchyme is a source of WNT, BMP and FGF ligands (Fig. 1a). We also observed that BTPs were the main source of TGFβ ligands in this analysis, which is predicted to signal in an autocrine manner as well as to non-BTP epithelium and SM22-positive mesenchyme (Fig. 1a). We have previously shown that activation of TGFβ- and BMP-signaling are associated with BTP differentiation into airway^23, 29^; therefore, to interrogate these pathways further, we analyzed predicted ligand-receptor pairs (Supplementary Fig. 1a) and plotted the expression of expressed ligands over developmental time in BTPs and RSPO2^+^ mesenchyme (Fig. 1b,c). This analysis revealed a trend of increasing transcription of BMP ligands, and the BMP-signaling target gene *ID2,* while TGFβ ligands remained unchanged or decreased over developmental time in both BTPs and RSPO2-positive mesenchyme. These changes correlated with decreasing levels of the BTP markers *SOX2* and *SOX9* as well as increased *SFTPC* expression in BTPs (Fig. 1b). To confirm increased BMP ligand and target gene expression with developmental time as suggested by our scRNA-seq analysis, we performed FISH for *BMP4* and *ID2* on tissue sections of pseudoglandular stage (59 days) and canalicular stage (112 days) human lung (Fig. 1d). Quantification of FISH foci in SOX9-positive BTPs, SOX9-negative (stalk, bud tip adjacent) non-BTP epithelium and their associated mesenchyme populations confirmed higher *BMP4* in BTPs (SOX9-positive) than non-BTP (SOX9-negative) cells at both timepoints examined, which significantly increased in both populations at the later timepoint (Fig. 1e). *ID2* transcription was also higher in BTPs compared to non-BTP epithelium at both timepoints and decreased in non-BTP epithelium at the later timepoint (Fig. 1f). These observations suggest that BTPs move towards a state of higher BMP- and lower TGFβ-signaling activity as development proceeds.

**Fig. 1:**
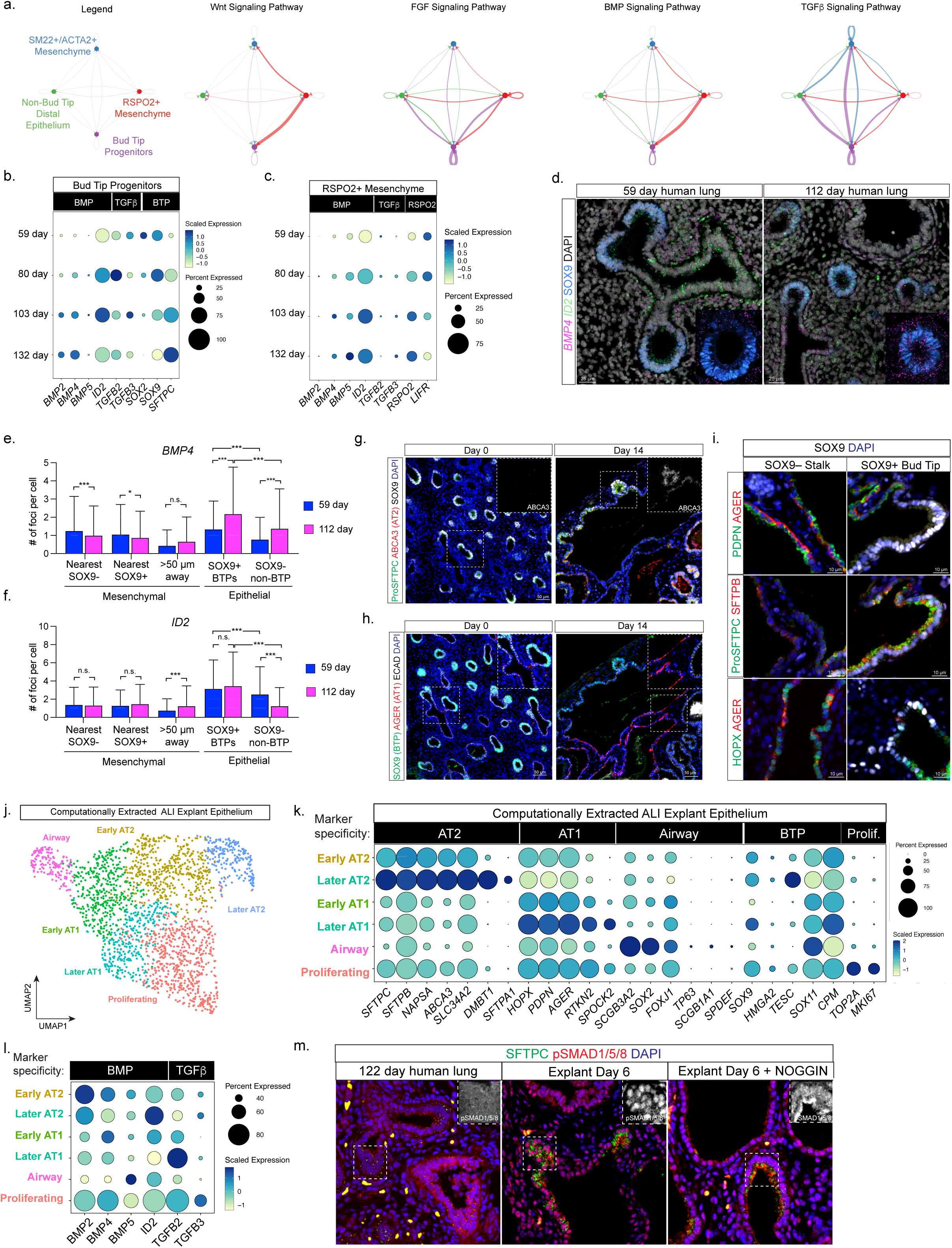
High levels of BMP- and low levels of TGFβ-signaling are associated with AT2 differentiation. a) Circle diagrams showing signaling network between cells in the lung progenitor niche (Bud Tip Progenitors, RSPO2+ mesenchyme) and cells outside (Differentiating Epithelium, TAGLN^+^ mesenchyme (myofibroblasts and smooth muscle)) for WNT-, FGF-, BMP- and TGFβ-signaling pathways. Interaction edges are colored by source signaling pathway and thickness represents the relative strength of the interaction. b) Dot plot showing expression of BMP and TGFβ ligands and BMP-signaling target *ID2* in BTPs. c) Dot plot showing expression of BMP and TGFβ ligands and BMP-signaling target *ID2* in RSPO2+ mesenchyme. d) Fluorescent *in situ* hybridization for *BMP4* and *ID2* with co-immunofluorescent staining for SOX9 in pseudoglandular (59 day) and canalicular (112 day) stage human lung. e) Quantification of *BMP4* FISH signal in mesenchymal and epithelial cell populations in distal lung sections at 59 and 112 days. Mesenchymal cells were classified based on their proximity to SOX9 expressing epithelial cells and epithelial cells were classified based on SOX9 expression. Statistical significance (p) was calculated by One-way ANOVA with Bonferroni correction. (* = p < 0 .05, *** = p < 0.001, n.s. = not significant (p > 0.05)). f) Quantification of *ID2* FISH signal in mesenchymal and epithelial cell populations in distal lung sections at 59 and 112 days. Mesenchymal cells were subdivided based on their proximity to SOX9 expressing epithelial cells and epithelial cells were subdivided based on SOX9 expression. Statistical significance (p) was calculated by One-way ANOVA with Bonferroni correction. (* = p < 0 .05, *** = p < 0.001, n.s. = not significant (p > 0.05)). Error bars = standard deviation. g) Immunofluorescent staining images of BTP marker SOX9 and AT2 markers (ProSFTPC, ABCA3) in lung explants before (day 0) and after ALI culture (day 14). Images are from a single biological replicate and representative of n = 4 biological replicates. h) Immunofluorescent staining images of BTP marker SOX9, AT1 marker AGER and epithelial marker ECAD in lung explants before (day 0) and after 14 days ALI explant culture. Images are from a single biological replicate and representative of n = 4 biological replicates. i) Immunofluorescent staining images in bud tip (SOX9-positive) and stalk (SOX9-negative) regions of lung epithelium in ALI explants at day 14 for AT2 (ProSFTPC, SFTPB), AT1 (PDPN, AGER) and HOPX (expressed in AT1 and AT2 cells in human). j) UMAP visualization of Louvain clustering of epithelial cells from day 3, day 6, day 9 and day 12 of ALI explant culture. Cluster identities were assigned based on marker expression in part k. Epithelial cells were extracted after annotation of cluster identities in the full dataset (Supplementary Fig. 1f, g). k) Dot plot showing expression of AT2, AT1, Airway, BTP and proliferating cell markers in Louvain clusters identified in UMAP in part j. l) Dot plot showing expression of BMP and TGFβ ligands and BMP-signaling target *ID2* in explant epithelial cell clusters. m) Immunofluorescent staining for pSMAD1/5/8 and ProSFTPC in 122 day fetal lung, day 6 ALI explants and day 6 ALI explants cultured in the presence of 100 ng/mL NOGGIN. Explant images are from a single biological replicate and representative of n = 3 biological replicates.

Interrogating later stages of human lung development when alveolar differentiation occurs is challenging due to a lack of access to tissue. To overcome this limitation, we turned to air-liquid interface (ALI) explant cultures, which allows continued development in serum-free, growth-factor free media. We explanted small distal fragments of 15 – 18.5 week canalicular stage human lung into ALI culture on polycarbonate filters floating on serum and growth factor free media^21, 23, 30^ (Supplementary Fig. 1b,c). The lung epithelium remained healthy and underwent changes in marker expression consistent with alveolar differentiation over the course of 14 days (Fig. 1g,h). In BTPs identified by SOX9 expression, we observed the onset of the AT2 cell marker ABCA3, which was not detected in BTPs prior to culture, as well as an increased intensity of ProSFTPC staining (Fig. 1g). In SOX9-negative cells, we saw expression of the AT1 marker AGER, which was limited in tissue prior to culture (Fig. 1h). IF analysis also revealed the onset of additional markers indicative of AT2 differentiation in bud tips, including SFTPC and SFTPB co-expression (Fig. 1I) and SFTPA, although frequency of SFTPA within the pool of ProSFTPC-positive cells was low relative to adult AT2s (Supplementary Fig. 1d,e). PDPN was co-expressed in cells staining positive for AGER (Fig. 1i). HOPX was detected in both bud tip and stalk regions, although to a lesser extent in bud tips (Fig. 1i). Thus, canalicular stage ALI explants undergo nascent alveolar differentiation as evidenced by the onset of AT1 and AT2 markers.

To expand these findings, we carried out scRNA-seq on a timecourse of ALI explants at day three, six, nine and twelve of culture. This analysis confirmed the presence of epithelial cells expressing markers of AT1 (*AQP4*, *AGER*), AT2 (*SFTPC*) or airway (*SOX2*) identity as well as additional populations of mesenchymal, immune, endothelial and mesothelial cells from all timepoints examined (Supplementary Fig. 1f,g). Extraction and re-clustering of epithelial cells identified a single cluster of airway cells, a cluster of proliferative cells, two clusters with AT1 marker expression and two clusters with AT2 marker expression (Fig. 1j,k). When compared to BTPs prior to culture, non-airway epithelial clusters in ALI explants had higher expression of *AGER* or *SFTPC* further supporting the conclusion that AT1 and AT2 differentiation occurs in ALI explant cultures (Supplementary Fig. 1i). Based on recent descriptions of AT1 and AT2 populations in adult and developing lungs^31–34^, we classified clusters as ‘early’ and ‘later’ in the differentiation process, with later AT1s distinguished by increased *RTKN2* and *SPOCK2* expression, and later AT2s distinguished by increased expression of *DNMBT1* and *SFTPA1* (Fig. 1k). Early and later designations within AT1 and AT2 clusters based on marker expression were not strongly correlated with time in culture (Supplementary Fig. 1j,k), indicating that differentiation is non-synchronous in explants. Nevertheless, comparing AT2 markers by scRNA-seq in BTPs, explant AT2s and adult AT2s confirmed broad AT2 marker onset relative to BTPs even when considered in bulk (Supplementary Fig. 1l), except for *SFTPA1*, which was infrequently expressed relative to adult AT2 cells, as expected from IF analysis (Supplementary Fig. 1d, e). Examination of BMP- and TGFβ-signaling ligands in explant epithelial populations suggested that cells differentiating towards AT1 transcribed higher levels of TGFβ ligands while cells differentiating into AT2s transcribed higher levels of BMP ligands and the BMP target *ID2* (Fig. 1l). Indeed, relative to canalicular stage human fetal lung tissue, ALI explants had higher levels of BMP-dependent phosphoSMAD1/5/8, particularly in cells undergoing AT2 differentiation as indicated by SFTPC co-staining (Fig. 1m). We further observed that phosphoSMAD1/5/8 could be blocked by the BMP inhibitor NOG in explants (Fig. 1m). These findings associate higher levels of BMP-signaling with cells undergoing AT2 differentiation and higher levels of TGFβ-signaling with cells undergoing AT1 differentiation in ALI explants.

### BMP- and TGFβ-signaling exhibit opposing activities on AT2 differentiation in explants and BTP organoids

Given data from tissue and explants suggesting roles for BMP- and TGFβ- signaling during alveolar cell differentiation (Fig. 1), we functionally interrogated the role of BMP- and TGFβ-signaling in serum and growth factor-free ALI explant cultures, focusing on the AT2 lineage. We used ProSFTPC-positive cells within ALI explants to identify nascent AT2 cells and used SFTPA co-expression as a readout for more advanced AT2 differentiation. SFTPA was undetected or limited to very low expression in a few cells in untreated ALI explants (Fig. 2a). Inhibition of BMP-signaling by addition of recombinant NOGGIN to the underlying media did not change SFTPA staining intensity or frequency relative to untreated ALI explants (Fig. 2a’). In contrast, activation of BMP-signaling with recombinant BMP4 led to robust and readily detectable levels of SFTPA staining in ProSFTPC-positive cells (Fig. 2a’’).

**Fig. 2:**
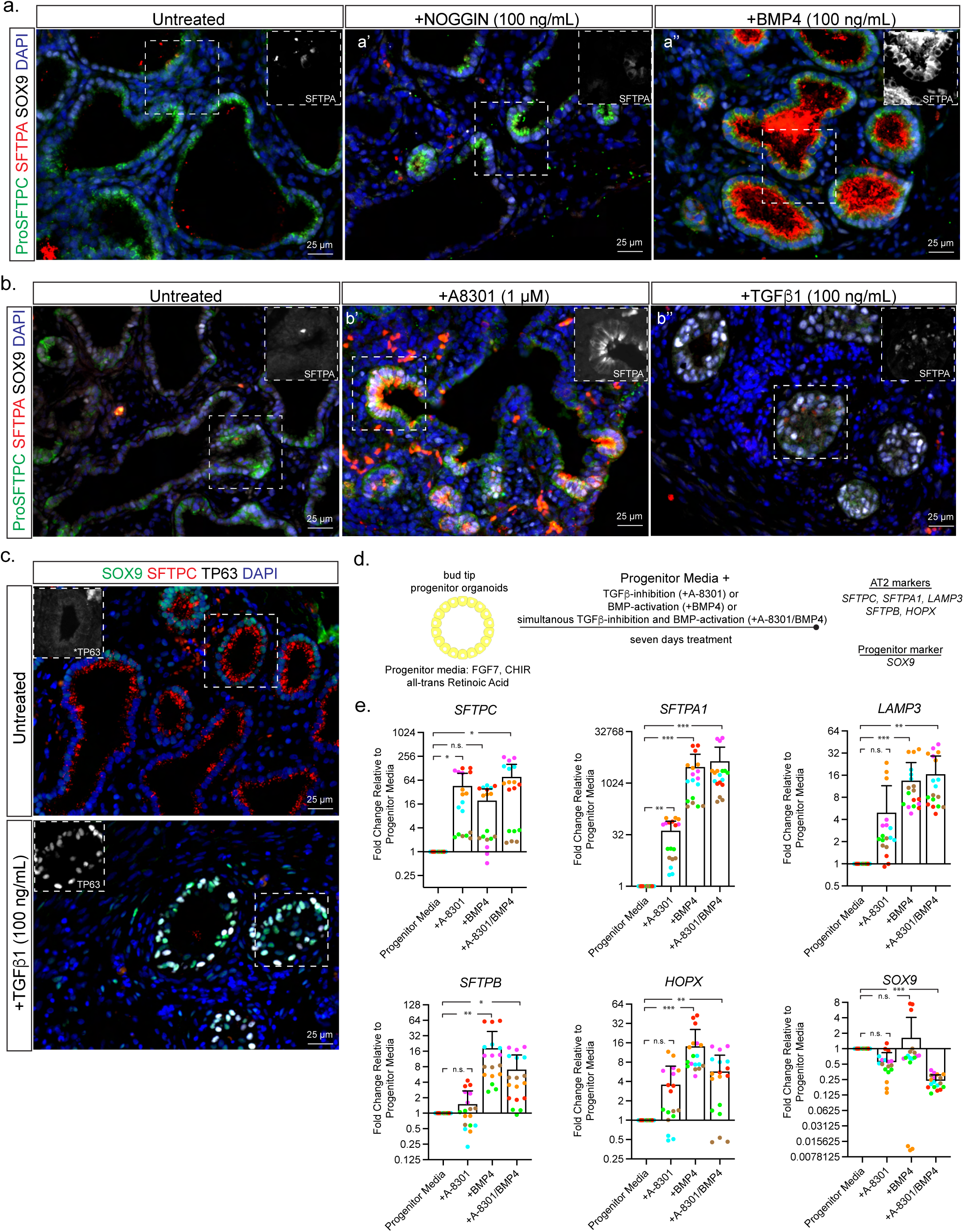
BMP- and TGFβ-signaling exhibit opposing activities on alveolar type 2 differentiation in explants and BTP organoids. a) Immunofluorescent staining images of AT2 marker (ProSFTPC and SFTPA) and BTP marker (SOX9) expression in (a’) BMP-inhibited (+NOGGIN) or (a’’) BMP-activated (+BMP4) day 6 ALI explant cultures of canalicular stage human lung showing increased SFTPA staining in response to BMP-activation. Images are from a single biological replicate and representative of n = 3 biological replicates. b) Immunofluorescent staining images of AT2 marker (ProSFTPC and SFTPA) and BTP marker (SOX9) expression in (b’) TGFβ inhibited (+A8301) or (b’’) TGFβ-activation (+TGFβ1) day 6 ALI explant cultures of canalicular stage human lung demonstrating increased SFTPA staining in response to TGFβ-inhibition. Images are from a single biological replicate are shown and representative of n = 3 biological replicates. c) Immunofluorescent staining images of BTP marker (SOX9), AT2 marker (ProSFTPC) and basal cell marker (TP63) expression in untreated and TGFβ1-treated explants showing ectopic TP63 in bud tip regions of TGFβ1-treated ALI explants. Images are representative of n = 3 biological replicates. d) Schematic describing approach to compare the effect of individual and simultaneous TGFβ- inhibition and BMP-activation on AT2 marker expression in BTP organoids by RT-qPCR. e) RT-qPCR measurements of AT2 markers *SFTPC*, *SFTPA1, LAMP3, SFTPB, HOPX* and BTP marker *SOX9* in BTP organoids cultured in progenitor media with addition of a TGFβ- inhibitor (+A8301), a BMP-activator (+BMP4) or both (+A8301/+BMP4) for seven days. Values shown are fold change relative to the progenitor media condition for each technical replicate color-coded by biological replicate. Statistical comparison (p) was calculated using repeated measures one-way ANOVA with Dunnett’s post-hoc test on linearized (log-transformed) mean fold-change values for each biological replicate calculated from the mean of technical replicates. (* = p < 0 .05, ** = p < 0.01, *** = p < 0.001, n.s. = p > 0.05). Error bars = standard deviation.

Manipulation of TGFβ-signaling had the opposite effects on SFTPA expression. Here, SFTPA was increased in ProSFTPC positve cells upon addition of the TGFβ-signaling inhibitor A-8301 (Fig. 2b’). In contrast, activation of TGFβ-signaling by the addition of recombinant TGFβ1 led to reduced ProSFTPC and barely detectible SFTPA staining (Fig. 2b’’), with most epithelial cells expressing TP63 (Fig. 2c), consistent with previous reports that TGFβ-activation promotes differentiation of BTPs towards airway^23, 35^. Taken together these results show that TGFβ-inhibition and BMP-activation led to the strongest increases in SFTPA expression, suggesting functionally opposing roles of TGFβ- and BMP-signaling in AT2 differentiation.

To corroborate the results from explant cultures we turned to epithelial only ‘BTP organoids’, in which the BTP state is maintained in progenitor media consisting of a WNT-agonist CHIR099021, FGF7 (otherwise known as Keratinocyte Growth Factor -KGF) and *all-trans* retinoic acid (ATRA)^3^. Given the AT2 promoting effects of BMP-activation or TGFβ-inhibition in explants, we hypothesized that activation of BMP-signaling or inhibition of TGFβ-signaling in BTP organoids would differentiate BTP organoids towards the AT2 lineage. In addition, we hypothesized that combining both signaling cues through simultaneous TGFβ-inhibition and BMP-activation would lead to more robust differentiation than manipulating each signaling pathway independently. To test this, we supplemented bud tip progenitor media with A-8301 or BMP4 individually, or A-8301 and BMP4 in combination for seven days and compared the response of AT2 differentiation markers by RT-qPCR (Fig. 2d).

Inhibition of TGFβ-signaling (+A-8301) significantly increased *SFTPC* and *SFTPA1* over levels observed in bud tip progenitor media (Fig. 2e). Activation of BMP-signaling (+BMP4) significantly increased *SFTPA1*, *LAMP3*, *SFTPB* and *HOPX* (Fig. 2e). Neither treatment alone significantly reduced expression of the BTP marker *SOX9*. However, exceeding the effects of individual treatments, simultaneous TGFβ-inhibition and BMP- activation (+A-8301/BMP4) significantly increased all AT2 markers examined while also significantly decreasing the BTP marker *SOX9* (Fig 2e). Examination of ProSFTPC, SFTPA and SOX9 protein by immunofluorescence corroborated the results of qPCR experiments, with +A-8301/BMP4 treated cells coexpressing ProSFTPC and SFTPA while SOX9 was decreased (Supplementary Fig. 2a). These experiments were performed in bud tip progenitor maintenance media containing FGF7, WNT-agonist CHIR90221, and ATRA. The FGF and WNT-signaling pathways are important components of the bud tip progenitor and AT2 niche *in vivo* and likewise are important for differentiation and maintenance of AT2s *in vitro*^2, 3, 5, 17, 21, 33, 36–38;^ however, the role of ATRA in AT2 differentiation is less clear^39, 40^. To test if ATRA is required for AT2 differentiation of BTPs we compared the effect of CHIR99021/KGF (‘CK’) plus A-8301/BMP4 (‘AB’) (referred to as CK + AB) in the presence or absence of ATRA by RT-qPCR. Removal of ATRA had mixed effects on AT2 differentiation, significantly enhancing *SFTPC* while reducing *SOX9,* while other markers were unchanged and *SFTPB* was reduced (Supplementary Fig. 2b). Because ATRA removal had very modest effects and did not positively or negatively impact AT2 differentiation we removed it in subsequent experiments.

### TGFβ-inhibition coupled with BMP-activation efficiently differentiates BTP organoids to AT2-like cells

To further evaluate the extent to which CK + AB is capable of differentiating BTP organoids towards an AT2 phenotype we further characterized cells differentiated with this method, focusing on the maturity of cells over the course of extended treatment, the efficiency of differentiation, and whether AT2-like cells made in this manner are committed to an AT2 identity when removed from differentiation media.

To examine the extent of AT2 differentiation in CK + AB treated BTP organoids, we first evaluated additional markers of AT2 identity by IF. Compared to cultures from the same passage maintained in bud tip progenitor media, we observed robust co-expression of ProSFTPC with additional AT2 markers including HTII-280^41^, NAPSA or SFTPA within 6 days of treatment (Fig. 3a). By RT-qPCR, CK + AB treated organoids increased AT2 markers over the course of 21 days and had reduced expression of the airway marker *SOX2* (Supplementary Fig. 3a). Expression of AT2 genes/proteins was maximal when BMP-signaling was activated, even if BMP-signaling was inhibited early during differentiation as BTP organoids cultured with A-8301 plus NOGGIN for 21 days had significantly less AT2 differentiation than parallel cultures in which BMP4 was added for the last seven days (Supplementary Fig. 3b-d). To determine if CK + AB treated cells possessed lamellar bodies, an important benchmark for AT2 maturation, we examined CK + AB treated BTP organoids at day 21 by Transmission Electron Microscopy (TEM). In contrast to the few and poorly formed lamellar bodies observed in spontaneously differentiated BTP organoids^3^, we observed abundant lamellar bodies located intracellularly and within the lumen of organoids (‘LB’ in Fig. 3b’). Moreover, lamellar bodies within the lumen appeared to be processed into tubular myelin, an ordered surfactant structure produced in the alveolar airspace^42, 43^ (Fig. 3b’’, 3b’’’), indicating AT2-like cells differentiated by CK + AB treatment possess functional capacity for lamellar body assembly, secretion, and extracellular processing.

**Fig. 3:**
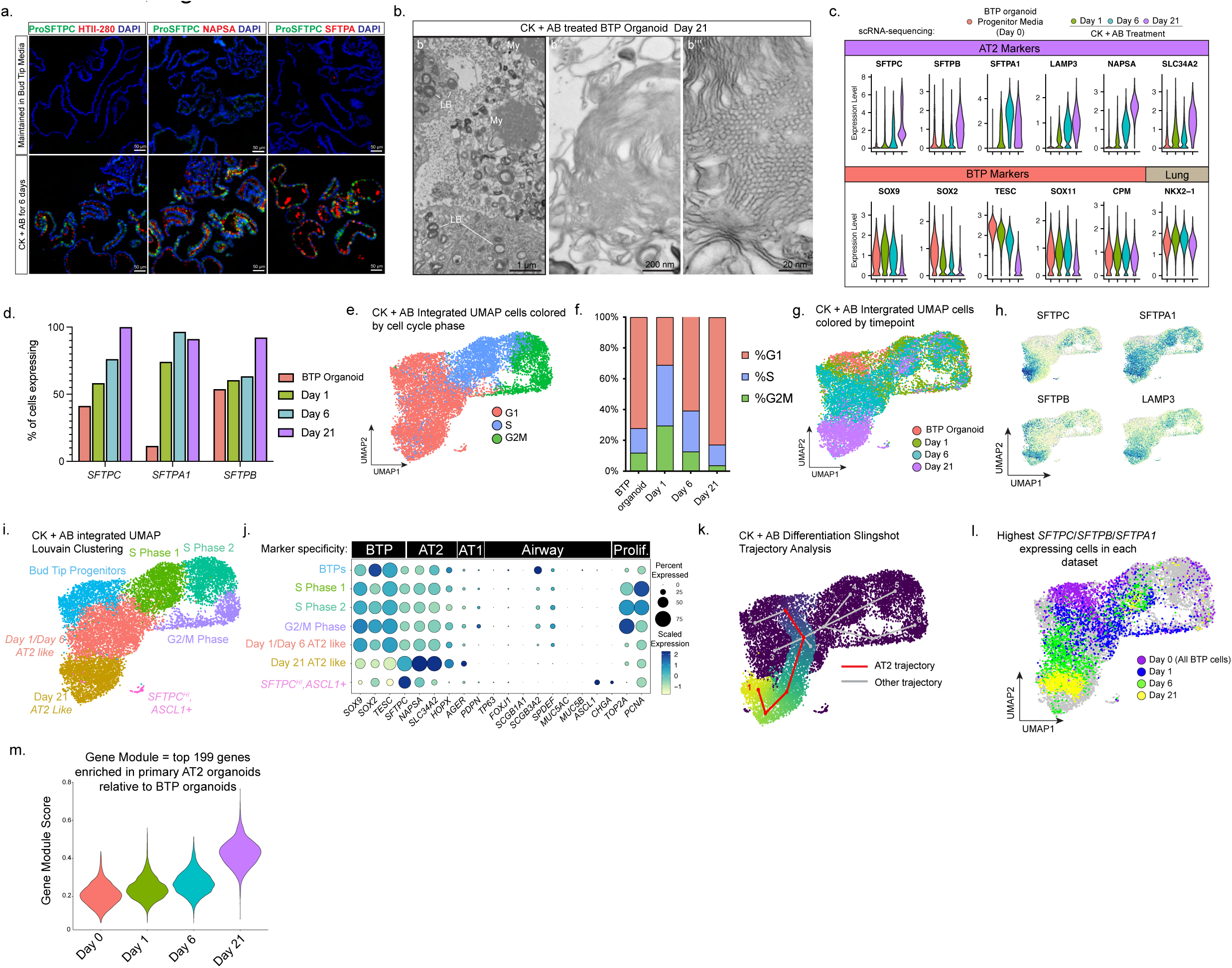
TGFβ-inhibition coupled with BMP-activation (CK + AB) efficiently differentiates BTP organoids to AT2-like cells. a) Immunofluorescent staining images of AT2 marker (ProSFTPC, HTII-280, NAPSA, SFTPA) onset and co-expression in BTP organoids treated for six days with CK + AB compared to cultures from the same passage maintained in bud tip progenitor media. b) Transmission electron microscopy images of lamellar bodies in BTP organoids after 21 days CK + AB treatment. c) Violin plots showing the distribution of AT2 and BTP marker gene expression from scRNA seq of untreated BTP organoids and at days 1, 6 and 21 of CK + AB treatment. d) Comparison of the proportion of cells expressing indicated AT2 marker in BTP organoids and at days 1, 6 and 21 of CK + AB treatment as determined by scRNA-seq. e) UMAP visualization of integrated scRNA-seq data from BTP organoids and day 1, 6 and 21 of CK + AB treatment showing with cells color coded by cell cycle stage. f) Comparison of the percentage of cells in each phase of the cell cycle in BTP organoids and at days 1, 6 and 21 of CK + AB treatment. g) UMAP visualization of integrated scRNA-seq data from BTP organoids and day 1, 6 and 21 of CK + AB treatment with cells color coded by sample origin. h) UMAP visualization of integrated scRNA-seq data from BTP organoids and day 1, 6 and 21 of CK + AB treatment with cells color coded by indicated AT2 marker gene expression. i) UMAP visualization of integrated scRNA-seq data from BTP organoids and day 1, 6 and 21 of CK + AB treatment with cells color coded by Louvain cluster. Cluster identities are labelled and were determined based on data shown in part e,f,g,h and j and cluster-specific enrichment lists. j) Dot plot showing expression (or absence) of key markers of proliferation, airway, AT1, AT2 and BTP identity across Louvain clusters from integrated CK + AB treatment scRNA-seq. k) Slingshot trajectory analysis of integrated scRNA-seq data from BTP organoids and day 1, 6 and 21 of CK + AB treatment. Trajectory originating in BTP organoids and terminating in area of day 21 high AT2 marker expressing cells is highlighted in red. Alternative trajectories are highlighted in gray and indexed for referencing in the main text. l) UMAP visualization of integrated scRNA-seq data from BTP organoids and day 1, 6 and 21 of CK + AB treatment with highest *SFTPC/SFTPB/SFTPA1* co-expressing cells from independent analysis of each scRNA-seq timepoint (Supplementary Fig. 3e-g) highlighted and color coded by timepoint. m) Violin plots comparing AT2 gene module scores for BTP organoids and day 1, 6 and 21 CK + AB treatment.

To investigate broader transcriptional changes in response to CK + AB treatment we performed scRNA-seq on BTP organoids in progenitor media (day 0), or after 1, 6 or 21 days of CK + AB treatment. Comparison of each timepoint confirmed the expected onset of AT2 markers and downregulation of BTP markers (Fig. 3c). Analysis of individual CK + AB treatment timepoints show that at day 1 and day 6 CK + AB treated cells expressed classical AT2 surfactant markers heterogeneously. By day 21 AT2 marker expression was more uniform with all cells expressing *SFTPC* and greater than 90% co-expressing *SFTPA1* and/or *SFTPB* (Fig. 3d, Supplementary Fig. 3e-g). The data also revealed a strong cell cycle dynamic in our timecourse data with an initial burst of cell proliferation at day 1 and reduced levels of proliferation in day 21 CK + AB treated cells relative to cells in BTP organoids (Fig. 3e, f). Integration of all treatment timepoints organized cells either as progressing through the cell cycle, or by increasing AT2 marker onset, which correlated with timepoint (Fig. 3e,g,h). Louvain clustering recognized day 0 BTPs as a single cluster, clustered day 1 and day 6 of CK + AB treatment together and separate from day 21 AT2-like cells (Fig. 3i). None of the CK + AB treated clusters identified expressed high levels of airway epithelial cell type markers, except for a very small number (< 0.05% of cells) of *SFTPC* expressing cells that co-expressed neuroendocrine markers (*SFTPC^HI^/ASCL1^+^;* Fig. 3j).

Trajectory analysis using Slingshot^44^ confirmed a trajectory originating from BTP organoids and terminating at day 21 cells expressing high levels of *SFTPC* (Fig. 3k, Trajectory 1). In general, this trajectory aligned with the cells in G1 phase of the cell cycle (Fig. 3e) and was in the location of cells with the highest levels of *SFTPC/SFTPA1/SFTPB,* identified during analysis of individual timepoints and overlaid onto the integrated UMAP embedding (Fig. 3l, Supplementary Fig. 3e-g). Other trajectories identified by this analysis led to clusters corresponding to stages of the cell cycle (Trajectories 4, 5 and 6), or terminated in areas of lower AT2 marker expression, possibly representing differentiation bottlenecks (Trajectories 2 and 3). To extend our characterization beyond a handful of AT2 and BTP marker genes, we performed scRNA-sequencing on primary adult AT2 organoids to define a list of the top 199 genes (Supplementary Table 3) enriched in primary AT2 organoids relative to BTP organoids (see methods, Fig. 5). We then used this list to assign a gene set module score^45^ to each cell in CK + AB treated BTP organoids. This analysis confirmed increasing acquisition of an AT2 molecular signature over 21 days CK + AB treatment (Fig. 3m). In total, these results show that with continued CK + AB treatment most cells acquire gene transcription consistent with AT2 differentiation without the co-differentiation of off-target cell types, suggesting that CK + AB efficiently differentiates BTPs to an AT2 identity.

### AT2-like cells induced by CK + AB expand in media optimized for primary AT2 organoids yet exhibit phenotypic instability

Although gene and protein expression of CK + AB-induced AT2-like cells was similar to primary AT2 cells *in vitro* (Fig. 3), we observed a decrease in growth of induced AT2-like cultures over the course of CK + AB treatment, leading to very little proliferation/growth by 21 days of differentiation (Fig. 3f, Fig. 4a). Primary AT2 cells have recently been shown to have the capacity to undergo extensive self-renewal in serum-free monoculture if given the appropriate growth cues^46, 47^. Therefore, we hypothesized that the continued growth of day 21 CK + AB induced AT2-like organoids requires specific AT2 growth conditions following acquisition of an AT2 identity.

**Fig. 4:**
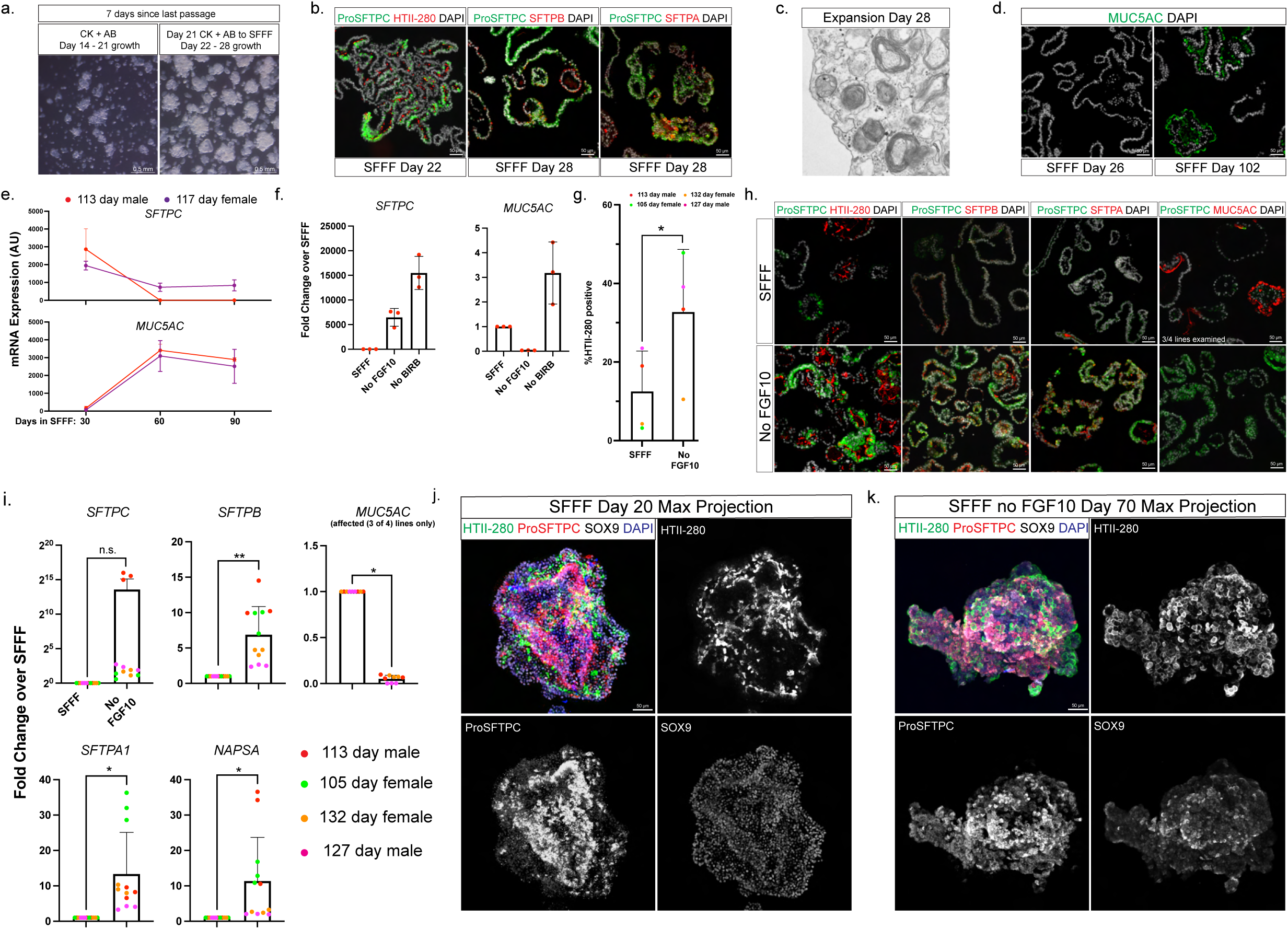
Optimization of expansion conditions for CK + AB induced AT2-like organoids. a) Brightfield images comparing growth between CK + AB induced AT2-like organoid cultures over days 14 – 21 in CK + AB media with the first 7 days in SFFF. b) Immunofluorescent staining images showing maintenance of ProSFTPC, HTII-280, SFTPB and SFTPA by CK + AB induced AT2-like organoid cultures early during expansion in SFFF. c) Transmission electron microscopy images of lamellar bodies in CK + AB induced AT2-like organoids after 28 days in SFFF media. d) Immunofluorescent staining images showing early absence and later appearance of MUC5AC-positive cells in CK + AB induced AT2-like organoids after expansion in SFFF media. e) RT-qPCR measurements of *SFTPC* and *MUC5AC* expression levels in CK + AB induced AT2-like organoids after 30, 60 and 90 days in SFFF media. Data is from differentiations of two BTP organoid lines with three technical replicates measured. f) RT-qPCR measurements of *SFTPC* and *MUC5AC* expression levels in complete SFFF or SFFF with indicated component (FGF10 or BIRB796) removed. Data is from a single BTP organoid differentiation with three technical replicates measured. g) Flow cytometry analysis of the proportion of HTII-280-positive cells in CK + AB induced AT2-like organoid cultures after 50 – 60 days in SFFF media with and without FGF10. Data is from differentiations performed on four independent BTP organoid lines. Statistical comparison (p) was computed by Student’s paired t-test. (* = p < 0 .05, ** = p < 0.01, *** = p < 0.001, n.s. = p > 0.05). Error bars = standard deviation. h) Immunofluorescent staining images comparing AT2 marker (ProSFTPC, HTII-280, SFTPB, SFTPA) and MUC5AC staining in CK + AB induced AT2-like organoid cultures after 90 days in SFFF media with and without FGF10. i) RT-qPCR measurements comparing AT2 marker (*SFTPC*, *SFTPB*, *SFTPA1*, *NAPSA*) and *MUC5AC* gene expression in CK + AB induced AT2-like organoids expanded after 90 days in SFFF with (SFFF) and without FGF10. Values shown are fold change relative to SFFF condition for each technical replicate color-coded by biological replicate. Statistical comparison (p) was computed by ratio paired t-test on the mean arbitrary units of expression for each biological replicate calculated from the mean of technical replicates. Expanded AT2-like organoids from the 105 day female line lost AT2 markers in the presence of FGF10 but did not upregulate *MUC5AC* and were excluded from statistical analysis for *MUC5AC*. (* = p < 0 .05, ** = p < 0.01, *** = p < 0.001, n.s. = p > 0.05). Error bars = standard deviation. j) Maximum signal intensity projection of confocal images from whole mount immunofluorescent staining for HTII-280, ProSFTPC and SOX9 in CK + AB induced AT2 like organoids showing domains of ProSFTPC-positive cells with inward facing HTII-280 and nuclear SOX9 at day 20 in SFFF media. k) Maximum signal intensity projection of confocal images from whole mount immunofluorescent staining for HTII-280, ProSFTPC and SOX9 in CK + AB induced AT2 like organoids showing domains of ProSFTPC-positive cells with HTII-280 facing outward from the organoid lumen and absence of nuclear SOX9.

**Fig. 5:**
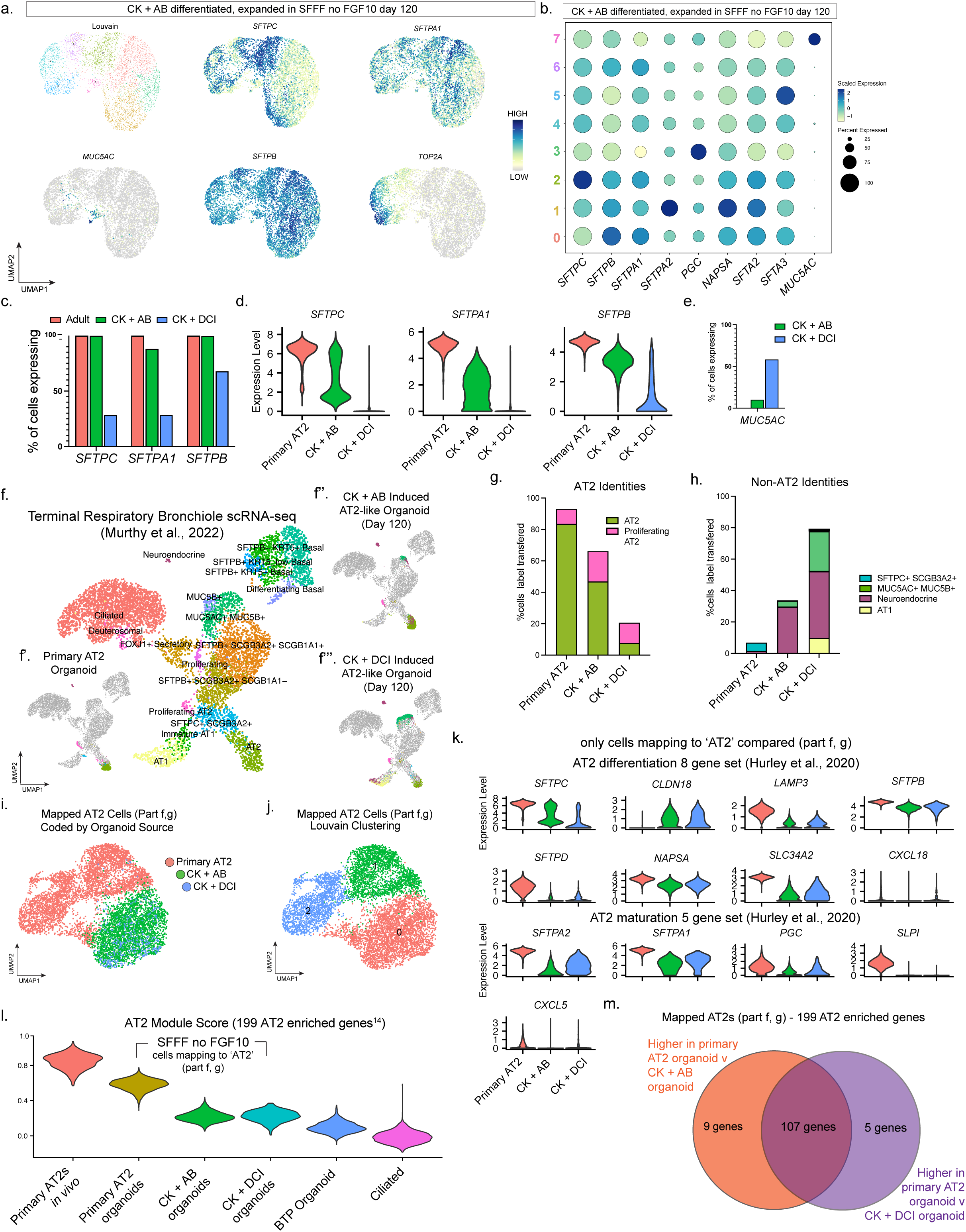
CK + AB differentiated organoids maintain AT2-like cells after long-term expansion. a) UMAP dimensional reduction with cells color coded by Louvain cluster or gene expression of AT2 (*SFTPC, SFTPA1, SFTPB)*, goblet (*MUC5AC*) and proliferation (*TOP2A)* markers in CK + AB induced organoids expanded for 120 days in SFFF without FGF10. b) Dot plot showing heterogenous but overall robust expression of an expanded list of AT2 marker genes and cluster 7 specific enrichment of goblet cell marker *MUC5AC* in CK + AB induced organoids expanded for 120 days in SFFF without FGF10. c) Comparison of the proportion of cells expressing indicated AT2 marker in primary AT2 organoids, CK + AB induced organoids and CK + DCI organoids after 120 days expansion in SFF without FGF10. d) Violin plots comparing the distribution of indicated AT2 marker expression levels in primary AT2 organoids, CK + AB induced organoids and CK + DCI organoids after 120 days expansion in SFFF without FGF10. e) Comparison of the proportion of cells expressing *MUC5AC* in CK + AB or CK + DCI induced organoids after 120 days expansion in SFFF without FGF10. f) Previously published UMAP dimensional reduction of proximal and distal lung scRNA sequencing. This data was used as a reference for unbiased mapping of cells from each indicated organoid type to known lung epithelial identities (f’ – f’’’). g) Percentage of cells mapping to AT2 identities from each indicated organoid type after 120 days expansion in SFFF without FGF10. h) Percentage of cells mapping to non-AT2 identities from each indicated organoid type after 120 days expansion in SFFF without FGF10. i) Cells mapping to the AT2 cluster in the reference from each organoid type were extracted, integrated and re-clustered. This UMAP dimensional reduction is color coded by organoid source to show AT2-like cells from induced organoids cluster together and separate from primary AT2 organoid cells. j) UMAP dimensional reduction of Louvain clustering for data in part i showing heterogeneity in primary AT2 organoid cells. k) Violin plots comparing gene expression levels in primary AT2, CK + AB and CK + DCI organoids for genes previously defined to characterize AT2 differentiation and maturation. Only cells mapping to AT2 cluster in the reference were compared. l) Violin plots showing gene module scores for the top 199 genes enriched in published scRNA sequencing data from human adult lungs^31^ (Supplementary Table 4) for indicated sample types. For organoid samples, only cells mapping to the AT2 cluster in the reference were compared, except for BTP organoids in which all cells from that dataset were analyzed. *In vivo* primary AT2s and Ciliated cells were extracted from the reference data set used in part f^48^. CK + AB and CK + DCI organoids score similarly, higher than BTP organoids and lower than primary AT2 organoids. m) Venn diagram showing overlap of AT2 marker genes (same gene set as module in part l) enriched in primary AT2 organoids relative to CK + AB or CK + DCI induced organoids. Only cells mapping to the AT2 cluster in the reference were compared. This comparison shows that despite the acquisition of an AT2 phenotype, the best AT2-like cells made with CK + AB or CK + DCI fall short of the gene expression levels seen in primary AT2 organoids for many of the same genes.

To test this, we transitioned day 21 CK + AB AT2-like organoids into serum-free feeder-free (SFFF) media optimized to support the self-renewal of primary AT2 cells in 3D organoid culture^46^. Day 21 CK + AB treated organoids transitioned to SFFF greatly increased their rate of growth relative to the amount of growth observed in day 14 – 21 CK + AB treated organoids (Fig. 4a) and exhibited extensive KI67 staining (Supplementary Fig. 4a), leading to highly proliferative cultures that could be passaged at a ratio of 1:6 every 7-14 days for at least 10 passages. Initially, AT2-like cells transitioned to SFFF were positive for AT2 markers ProSFTPC, HTII-280, SFTPB and SFTPA1 (Fig. 4b), and possessed lamellar bodies (Fig. 4c). With continued culture in SFFF, we observed loss of *SFTPC* expression and increasing expression of *MUC5AC* (Fig. 4d,e). This data indicates that the AT2 phenotype of CK + AB-induced cells is unstable in SFFF.

### Removal of FGF10 from SFFF reduces phenotypic instability of CK + AB-induced AT2 cells

We hypothesized that CK + AB-induced AT2 cells may be sensitive to growth factors or other components in SFFF, given that the expansion media was developed and optimized for fully mature AT2 cells from adults. We tested two modified versions of SFFF, one without the p38 MAPK inhibitor BIRB797, and the other without FGF10. Interestingly, removal of BIRB797 or FGF10 led to improved expression of *SFTPC* (Fig. 4f). Additionally, removal of FGF10 led to a reduction of *MUC5AC* to near-zero levels, while removal of BIRB797 increased *MUC5AC* (Fig. 4f). Based on this data, additional experiments were carried out to compare the robustness of SFFF without FGF10 to maintain the AT2 phenotype of CK + AB induced AT2-like cells.

CK + AB induced AT2-like cells from multiple biological specimens were transitioned to SFFF with and without FGF10 for 60 days and interrogated by FACS to determine the percent of cells expressing HTII-280 (Fig. 4g), by IF for co-expression of AT2 markers (ProSFTPC, HTII-280, SFTPB, SFTPA) and MUC5AC expression (Fig. 4h) and by qRT-PCR to measure bulk expression levels of these markers (Fig. 4i). This data confirmed the robustness of SFFF without FGF10 to maintain AT2 gene/protein expression while expanding CK + AB induced AT2-like organoids.

We noted differences in the appearance of induced AT2-like cells/organoids grown in SFFF with or without FGF10. Organoids maintained in SFFF with FGF10 primarily possessed a cystic appearance and localized HTII-280 on the luminal surface of organoids (Fig. 4j, Supplementary Fig. 4b). In contrast, organoids maintained in SFFF without FGF10 grew as clusters of cells with HTII-280 on the membrane facing away from the organoid lumen (Fig. 4k, Supplementary Fig. 4c), similar to primary AT2 cultures in complete SFFF (Supplementary Fig. 4d). CK + AB-induced AT2-like organoids maintained many SFTPA-positive cells that colocalized with ProSFTPC (Supplementary Fig. 4e). Collectively, these results indicate that the AT2 phenotype of CK + AB differentiated AT2-like cells is repressed by FGF10, and that AT2-like cells expanded in SFFF without FGF10 maintain AT2 marker expression.

### CK + AB induced organoids maintain AT2-like cells after long-term expansion

To determine the similarity of expanded CK + AB induced AT2 organoids to primary AT2 organoids we carried out a direct head-to-head comparison in SFFF without FGF10 media by scRNA-seq. Additionally, given that well established methods have been developed to generate iPSC-derived AT2 cells, we also included this method in the comparison by treating BTP organoids with CK + DCI as previously described^5, 15–17^. BTP organoids were differentiated with either CK + AB or CK + DCI for 21 days and then expanded for an additional 120 days in SFFF without FGF10. Primary AT2 organoids were established from HTII-280^-positive^ cells isolated from adult (>60 years, deceased) distal lung cultures, expanded in SFFF for 90 days and passaged into SFFF without FGF10 media 30 days before scRNA-sequencing.

We first evaluated each dataset independently to assess the proportion of cells exhibiting an AT2 phenotype and to identify non-AT2 cell types (Fig. 5). Within CK + AB induced AT2 cells, heterogeneity was observed and highlighted by cluster specific enrichment of individual AT2-associated genes such as *SFTPC* and *SFTPA1*, whereas some genes such as *SFTPB* were uniformly expressed across all cells (Fig. 5a,b). In contrast, in primary AT2 cultures expression of *SFTPC*, *SFTPA1* and *SFTPB* was uniformly high (Supplementary Fig. 5a,b). When compared to CK + DCI (Supplementary Fig. 5c,d), AT2-like organoids differentiated with CK + AB had a greater proportion of cells expressing (Fig. 5c) and higher expression levels of genes encoding surfactant proteins (Fig. 5d). Both CK + AB and CK + DCI induced organoids contained *MUC5AC*^-positive^ cells, suggestive of a goblet cell identity, although to a much greater extent in CK + DCI induced organoids (Fig. 5e). Increased *MUC5AC* expression in expanded CK + DCI induced organoids relative to CK + AB organoids was reproducible across differentiations performed on multiple BTP organoid lines (Supplementary Fig. 5e). *MUC5AC*-positive cells appeared to be the main off-target cell type in induced organoids, as markers of other cell types were either not broadly expressed or lower in induced relative to primary AT2 organoids (Supplementary Fig. 5f). Gene module scoring of *MUC5AC*-positive cells in CK + DCI and CK + AB using gene enrichment lists from two goblet cell populations in an adult reference dataset^48^ suggested that *MUC5AC positive* goblet-like cells were immature in both conditions (Supplementary Fig. 5g,h). Based on these observations, we conclude that CK + AB AT2-like organoids maintain a larger proportion of cells with AT2-phenotype than CK + DCI AT2-like organoids in SFFF without FGF10.

To support this conclusion, we performed “referenced-based” mapping^49^ of all three organoid types using scRNA-seq of adult human terminal airways and alveoli^48^ as a reference (Fig. 5f). The majority of cells from primary AT2 organoids mapped to AT2 or proliferating AT2 clusters (93%), with smaller proportions mapping to a recently discovered cell type in terminal respiratory bronchioles co-expressing *SFTPC* and *SCGB3A2*^48^ (5%) or the neuroendocrine (1%) cluster (Fig. 5f’,g, h). In day 120 CK + AB induced AT2-like organoids the majority (66%) of cells mapped to either AT2 or proliferating AT2 cells (Fig. 5f’’, g), with the remainder of the cells mapping to either neuroendocrine cells (29%) or MUC5AC^+^MUC5B^+^ goblet cells (4%) (Fig. 5f’’,h). In contrast to primary or CK + AB induced organoids, most cells in day 120 CK + DCI induced AT2-like organoids mapped to non-AT2 cell types, which comprised neuroendocrine cells (42%), MUC5AC^+^MUC5B^+^ goblet cells (25%) and AT1 cells (10%), leaving a smaller fraction of cells mapping to AT2 or proliferating AT2 cells (21%) (Fig. 5f’’’,g, h). These results are consistent with our clustering and analysis of cell identity markers and indicate that treatment of BTP organoids with CK + AB generates a more homogenous population of AT2 cells than CK + DCI when expanded in SFFF without FGF10 reflecting commitment of CK + AB AT2-like cells towards an AT2 phenotype.

### Expanded AT2-like cells in CK + AB and CK + DCI organoids are transcriptionally similar to each other and distinct from primary AT2 organoids

To assess the degree of similarity between cells retaining AT2-like cells in all three types of organoids, we extracted CK + AB and CK + DCI treated cells mapping to AT2 clusters in the adult human reference dataset (Fig. 5f) and used batch correction to integrate them along with primary AT2 organoid cells into a common UMAP (Fig. 5i,j). CK + AB and CK + DCI induced AT2-like cells clustered together, with little mixing into primary AT2 organoid cells (Fig. 5i), indicating that AT2-like cells differentiated with CK + AB or CK + DCI are transcriptionally similar to each other and distinct from primary AT2 organoids. In fact, lower resolution Louvain clustering revealed that AT2 mapping cells from primary AT2 organoids have more transcriptional heterogeneity amongst themselves than CK + AB or CK + DCI mapped AT2 cells have between each other (Fig. 5j). These findings are consistent with prior reports that AT2s differentiated *in vitro* retain an immature phenotype relative to primary AT2 organoids^50^.

Comparing expression of AT2 marker genes previously reported to describe AT2 differentiation and maturation^25^, we noted that expression of many of these genes was similar between AT2 mapping cells in induced organoids but was lower relative to cells from primary AT2 organoids (Fig. 5k). To compare a broader set of AT2 markers we utilized a previously published list of 199 genes enriched in AT2 cells relative to all other lung cell types^31^ to calculate a module score^45^ for every cell mapping to the AT2 cluster in the reference dataset. As controls, we also included scRNA-seq data from *in vivo* primary AT2s^48^, and non-AT2 cell types including primary multiciliated cells^48^ and BTP organoids (this study). This analysis (Fig. 5i) scored cells retaining an AT2 phenotype from both differentiation methods similarly, and lower than primary AT2 organoids, supporting our conclusion from examining a smaller targeted list of AT2 marker genes (Fig. 5k). In addition, differential expression analysis revealed extensive overlap in AT2 markers expressed higher in primary AT2 organoids than AT2 mapping cells from both types of induced AT2-like organoids (Fig. 5m).

Based on this comparison between the ‘best’ AT2-like cells from CK + AB, CK + DCI induced and primary AT2 organoids we conclude that under the expansion conditions used here, AT2-like cells induced by both methods possess an equivalent AT2 phenotype and are characterized by expression of many AT2 genes, but at lower levels relative to primary AT2 organoids.

### CK + DCI derived AT2 cells transition through an *SCGB3A2*-positive intermediate state not found in CK + AB differentiation

To further interrogate differences between CK + AB and CK + DCI induced AT2-like organoids, we performed a scRNA-seq timecourse of cultures from day 1, 6 and 21 of CK + DCI differentiation (Fig. 6) to time-match the data we generated for CK + AB induction (Fig. 3). Similar to results from CK + AB differentiations, we saw the onset of AT2 marker expression throughout most of the culture by day 21 (Fig. 6a). Cell cycle activity decreased upon CK + DCI treatment, rather than increased as observed CK + AB time-series data (Supplementary Fig. 6a, Fig 3f). Integration of this time-series with undifferentiated ‘Day 0’ BTP organoids arranged day 1 and day 6 samples so that they were overlapping, and distinct from day 21 or day 0 cells (Fig. 6b-d). Clusters primarily comprised of cells from day 21 CK + DCI were present expressing markers of basal (*TP63*) and early-neuroendocrine identity (*ASCL1*), although relatively minor in size relative to the day 21 AT2-like cluster (Fig. 6b-d). Comparison between the *ASCL1^+^* clusters in CK + AB (Fig. 3) and the CK + DCI differentiations revealed statistically higher levels of ASCL1 in CK + DCI cells and similar levels of AT2 (*SFTPC*) or mature neuroendocrine (*CHGA*) markers (Supplementary Fig. 6b). This analysis also identified two additional CK + DCI clusters from day 1 and day 6 cells not found in CK + AB differentiations that were defined by high solute transporter expression (Unknown 1) or unique expression of *CXCL8, TF* and *LCN2* (Unknown 2) (Supplementary Fig. 6c). Thus, similar to CK + AB differentiations, the vast majority of cells differentiate towards AT2 identity in response to CK + DCI treatment, although CK + DCI contains additional rare sub populations not found in CK + AB, including unique clusters of day 1 and 6 cells as well as day 21 basal-like cells.

**Fig. 6:**
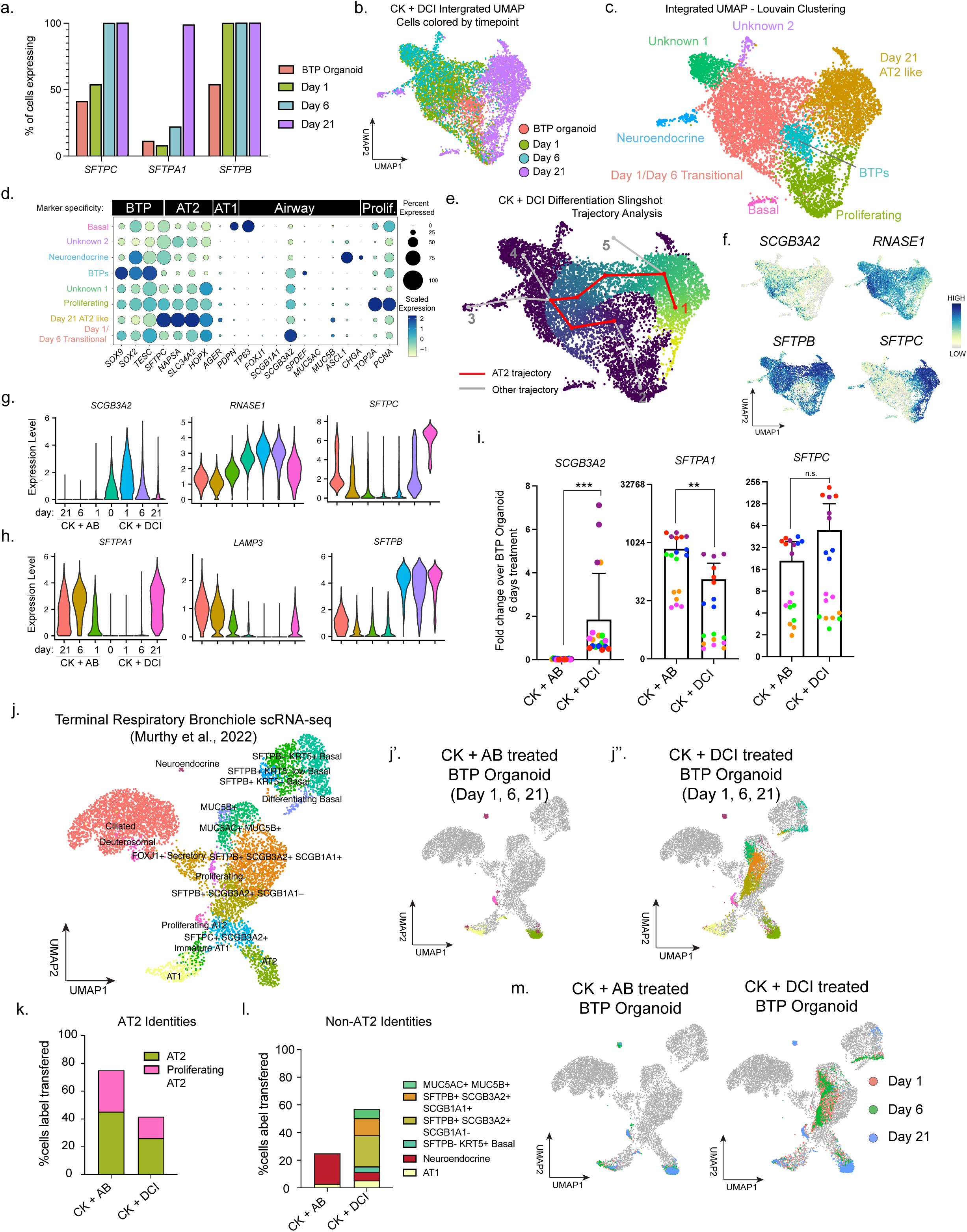
CK + DCI derived AT2 cells transition through an *SCGB3A2*-positive intermediate state not observed in CK + AB differentiation. a) Comparison of the proportion of cells expressing indicated AT2 marker in BTP organoids and at days 1, 6 and 21 of CK + DCI treatment as determined by scRNA-seq. b) UMAP dimensional reduction of integrated scRNA-seq data from BTP organoids and day 1, 6 and 21 of CK + DCI treatment with cells color coded by sample origin. c) UMAP dimensional reduction of integrated scRNA-seq data from BTP organoids and day 1, 6 and 21 of CK + DCI treatment with cells color coded by Louvain cluster. Cluster identities are labelled and were determined based on data shown in part b and f and examination of cluster specific enrichment lists. d) Dot plot showing expression (or absence) of key markers of proliferation, airway, AT1, AT2 and BTP identity across Louvain clusters from integrated CK + DCI treatment scRNA-seq data. e) Slingshot trajectory analysis of integrated scRNA-seq data from BTP organoids and day 1, 6 and 21 of CK + DCI treatment. Trajectory originating in BTP organoids and terminating in area of day 21 high AT2 marker expressing cells are highlighted in red. Alternative trajectories are highlighted in gray and indexed for referencing in the main text. f) UMAP visualization showing gene expression for markers enriched at Slingshot trajectory branching point and AT2 marker *SFTPC*. g) Violin plots of gene expression showing differential expression of indicated markers in BTP organoids responding to CK + AB or CK + DCI. h) Violin plots of gene expression showing differential expression of AT2 markers in BTP organoids responding to CK + AB or CK + DCI. i) RT-qPCR measurements of gene expression at day 6 relative to starting levels in BTP organoids showing increased *SCGB3A2* in CK + DCI, increased *SFTPA1* in CK + AB and similar levels of *SFTPC* across multiple BTP organoid lines. j) Previously published UMAP dimensional reduction of proximal and distal lung scRNA-sequencing. This data was used as a reference for referenced based mapping of cells from day 1, 6 and 21 of BTP organoids treated with CK + AB or CK + DCI (j’ – j’’). CK + DCI treated cultures contain cells mapping to terminal respiratory bronchiole identities. Similar cells are not seen in CK + AB differentiations. k) Percentage of cells treated with CK + AB or CK + DCI mapping to AT2 identities (part j). l) Percentage of cells treated with CK + AB or CK + DCI mapping to non-AT2 identities (part j). m) UMAP visualization of query datasets projected onto reference dataset and color coded by day of treatment. This visualization shows that cells mapping to terminal respiratory bronchiole identities come primarily from day 1 and day 6 of CK + DCI treatment.

To ascertain the trajectory of cells differentiating towards AT2 identity in response to CK + DCI, we performed Slingshot analysis on the integrated time-series data and found a trajectory transitioning through day 1 and 6 cells and terminating in day 21 AT2-like cells (Fig. 6e, Trajectory 1). This trajectory was in general agreement with the location of cells expressing the most *SFTPC/SFTPA1* identified during analysis of individual timepoints and overlaid on the integrated UMAP embedding (Supplementary Fig. 6d-g). We also observed a trajectory corresponding to proliferation (Trajectory 2) and additional trajectories branching off from the AT2 trajectory to terminate in areas of neuroendocrine marker (Trajectory 3) or clusters of unknown identity from day 1/6 (Trajectory 4) (Fig. 6e). Examining gene expression at this major branching point, we observed *SCGB3A2 and RNASE1* within the top 10 enriched genes. Expression of *SCGB3A2* and *RNASE1* preceded the onset of *SFTPC* along the AT2 trajectory (Fig. 6f), suggesting *SCGB3A2/RNASE1* expression marks cells differentiating to an AT2 identity in response to CK + DCI. Examination of gene expression during CK + AB differentiations showed that in contrast to CK + DCI, CK + AB treatment leads to sustained downregulation of *SCGB3A2* and *RNASE1*, and similar pace of *SFTPC* onset (Fig. 6g). CK + AB response is further distinguished by earlier onset of *SFTPA1* and *LAMP3* and delayed onset of *SFTPB* relative to CK + DCI (Fig. 6h). These observations were reproducible by RT-qPCR across differentiations from multiple BTP organoid lines (Fig. 6i). Transcriptional differences between CK + AB and CK + DCI differentiations suggests differences in the transcriptional trajectory of cells as they acquire their AT2 identity in either differentiation media.

To understand the relevance of transcriptional differences between BTP organoid cells treated with CK + AB or CK + DCI to the alveolar region of the lung we again turned to reference based mapping to distal adult lung epithelium scRNA-seq data^48^ (Fig. 6j-l). Most cells from CK + AB differentiation mapped to AT2 identities, with the remainder mapping to AT1 and neuroendocrine identities (Fig. 6j’,k,l). Cells from CK + DCI differentiation also contained many cells mapping to AT2, AT1 and neuroendocrine identities, but additionally mapped to basal, goblet, and both clusters of *SFTPB^+^/SCGB3A2^+^*secretory-like cells, which in this reference dataset represent epithelial cells specific to terminal respiratory bronchioles^37, 48^ (Fig. 6j’’,k, l). Examination of cells by day of differentiation revealed that cells from CK + DCI differentiations mapping to the *SFTPB^+^/SCGB3A2^+^* cluster in the reference were transitional, existing at days 1 and 6 but not detected at day 21 (Fig. 6m).

This comparison demonstrates that AT2-like cells induced by either CK + AB and CK + DCI acquire AT2 marker expression through distinct transcriptional trajectories, with CK + DCI expressing markers of terminal respiratory bronchiole identities (*SCGB3A2*/*RNASE1*) *en route* to AT2 cell identity.

## Discussion

Here we focused on the roles of TGFβ- and BMP-signaling during differentiation of human BTPs to AT2 cells. We show these pathways regulate AT2 differentiation in opposing fashion, with TGFβ-signaling acting to inhibit and BMP-signaling acting to promote AT2 differentiation. These activities are consistent with the proximal distal patterning of these signaling pathways during mouse lung development, with TGFβ-ligands restricted to proximal areas of the lung that comprise the future airways^51, 52^, and BMP-ligands restricted to bud tip progenitors in the distal areas of the lung that comprise the future alveoli^53^. Disruption of this patterning, either by overexpression of TGFβ1 in BTPs^54^, or overexpression of BMP inhibitors in BTPs^55, 56^ arrests lung development, emphasizing the importance of TGFβ and BMP patterning for proper lung development. Taken together our analysis of TGFβ- and BMP-signaling in the developing human lung as well as our functional experiments in ALI explant culture and lung experiments support a model where low levels of TGFβ-signaling in the distal lung and increasing levels of BMP-signaling combine with the high WNT- and FGF-signaling environment in the bud tip niche to promote the differentiation of BTPs to AT2 cells.

In addition to the roles described above, TGFβ-signaling is also required for branching morphogenesis^54, 57, 58^, airway homeostasis and regeneration^59, 60^, AT1 cell differentiation^61–63^, and BMP-signaling additionally regulates post-natal alveologenesis and AT2 cell homeostasis^64, 65^. Moreover, aberrant TGFβ-signaling has been proposed to contribute to many lung diseases, including bronchopulmonary dysplasia^66, 67^, idiopathic pulmonary fibrosis^68, 69^ and asthma^70, 71^. Thus, mechanisms regulating TGFβ- and BMP-signaling are of interest. In development, differential localization of specific mesenchymal populations, such as distally localized RSPO2+ mesenchyme, which we show here is a source of BMP ligand, or more proximally localized smooth-muscle and myofibroblast populations help to establish patterning. We also show here that canicular stage bud tips as late as 17.5 weeks (122 days) respond to TGFβ-signaling by upregulating airway differentiation markers, revealing that canicular stage bud tips remain competent for airway differentiation and suggesting that failure to repress TGFβ in airways would have catastrophic effects of bronchioalveolar organization of the lung epithelium, akin to lesions described in multiple subtypes of congenital lung malformations^72, 73^. Additionally, our analysis of pseudoglandular and canicular stage human lung shows that BTPs themselves contribute to the dynamics of TGFβ and BMP ligand availability, emphasizing an important contribution of BTPs to shaping their own niche.

TGF and BMP ligands are part of a larger family of ancestrally related Transforming Growth Factors that regulate stem cells through opposing and cooperative activities in many tissues^74–77^. Canonically TGFβ- and BMP-signaling use different receptor complexes, intracellular mediators, and transcriptional co-factors which converge on the DNA-binding protein SMAD4^78^. Work from our lab has shown that in the context of high TGFβ- signaling, BMP-signaling acts cooperatively to enhance airway differentiation of BTP organoids. Here we show that in the context of low TGFβ-signaling activity, BMP-signaling activity instead promotes AT2 differentiation of BTP organoids. Taken together these studies suggest that the balance between TGFβ- and BMP-signaling is a major determinant of cell fate in BTPs. This relationship mirrors that of studies in other organs where the balance of TGFβ- and BMP-signaling determines cell fate outcomes, with competition between TGFβ and BMP specific transcriptional co-factors for SMAD4 mediating crosstalk between signaling pathways^79–81^. Crosstalk between TGFβ- and BMP-signaling also occurs through protein-protein interactions and secondary messengers, which may be important for maintaining proximal-distal gradients of these pathways in the lung^82^. Interactions with other cell signaling pathways in the BTP niche like WNT- and FGF-signaling enhance the complexity of organ patterning and cell specification^83^. Airway and AT2 differentiation of BTP organoids provides a tractable model to further investigate mechanisms that translate TGFβ- and BMP-signaling levels into specific cell fates during human lung development.

Given that CK + AB and CK + DCI both induce the differentiation of BTP organoids to AT2-like cells an important question is whether both differentiation methods converge on similar mechanisms to accomplish AT2 differentiation. Glucocorticoids like dexamethasone have been reported to repress TGFβ-signaling in the lung^84, 85^, which would support the idea that both methods induce a similar signaling environment in BTPs. However, our scRNA-seq timecourse and of both differentiations shows that CK + DCI contain an SCGB3A2+ population not observed in CK + AB differentiations. The relevance of this gene expression difference between differentiation methods is not clear, but cells with *SFTPB+/SCGB3A2+* co-expression have been identified during development^23, 35^, as well as within terminal respiratory bronchioles of adults^37, 48^, suggesting that CK + DCI induced BTP organoids transit through a transcriptional state similar to these *in vivo* cell types prior to acquiring maximal AT2 identity. Consistent with this interpretation we find by referenced-based mapping that cells treated with CK + DCI align to SCGB3A2+/SFTPB+ cells in terminal respiratory bronchioles specifically at days 1 and 6. The absence of a similar population in CK + AB differentiations argues for differences in the mechanisms by which CK + AB and CK + DCI induce AT2 differentiation. Combining the *in vitro* models of human lung development described here and in the literature with gene knockout/down^86^ and lineage tracing approaches^35^ will be important to further interrogate the gene networks and transitional states required for human AT2 differentiation.

We noted AT2-like organoids generated with either CK + AB or CK + DCI become contaminated with *MUC5AC*-positive goblet-like cells over time, a process that is accelerated by the presence of FGF10. This mirrors results of FGF10 overexpression in the mouse lung, which results in goblet cell metaplasia within alveoli^87^. Recently multiple studies have converged on the presence of mislocalized airway basal cells in the distal lungs associated with Idiopathic Pulmonary Fibrosis (IPF)^88–90^, which are proposed to arise through pathological transdifferentiation events originating from AT2 cells. In CK + AB cultures markers of basal (*TP63*) and goblet (*SPDEF*, *MUC5AC*, *MUC5B*) cells are not detected at the timepoints sampled during 21 days of differentiation, arguing that the MUC5AC-positive goblet-like cells arising in SFFF originate from AT2-like after day 21. Likewise, we do not observe a robust population of TP63-positive cells in either CK + AB or CK + DCI cultures after long-term expansion in SFFF without FGF10, suggesting *MUC5AC* expressing cells arise without a basal cell intermediate state. Thus, the AT2-like to goblet-like transition observed here seems distinct from what has been reported in IPF patients, where basal cells are present. Never-the-less goblet cell metaplasia is associated with IPF and other lung diseases including chronic obstructive pulmonary disorder (COPD), pulmonary infections and cancer^91–95^, making the AT2-to-goblet cell fate transition observed here worthy of deeper investigation.

While we were able to investigate nascent alveolar cell differentiation using ALI explants of canicular stage lung, this study is limited by lack of access to human tissue specimens from later stages of lung development when *bona fide* alveolar differentiation is occurring. Our results show that ALI cultures initiate alveolar cell differentiation, and explants facilitate gain- and loss-of-function experiments and are therefore useful for probing the cell signaling pathways that regulate AT1 and AT2 differentiation; however, how closely ALI explants model later stage AT1 and AT2 differentiation is not known. Additionally, access to human tissue specimens at later stages would permit alignment of the transcriptional trajectories identified in AT2 induction using scRNA-sequencing timecourse data (i.e. CK + AB or CK + DCI differentiation) to *in vivo* development. Given the fact that we identified different trajectories for each of these differentiation methods, aligning these trajectories with *in vivo* differentiation would help reveal the relevance of these different trajectories to human alveolar development, homeostasis and regeneration.

Taken together, our data presented herein shows that TGFβ- and BMP-signaling work in opposition to regulate AT2 differentiation of BTPs during lung development. Translating these observations to *in vitro* organoid cultures, we show that inducing a state of low TGFβ- and high BMP-signaling activity, when combined with high WNT- and FGF-signaling activity, leads to robust differentiation of AT2-like organoids from BTP organoids *in vitro*. We anticipate AT2-like cells generated with CK + AB to complement existing methods to generate AT2-like cells, providing a valuable model for investigation of human lung biology and regeneration.

## Methods

### Experimental Model and Subject Details

#### Human lung tissue

All research utilizing human lung tissue (8 – 18.5 weeks post conception lung, adult lung) was approved by the University of Michigan Institutional Review Board. Human fetal lung tissue specimens were from presumably normal, de-identified specimens processed by the University of Washington Laboratory of Developmental Biology. Specimens included both male and female sexes. Tissue was shipped in Belzer-UW Cold Storage Solution (Thermo Fisher, Cat#NC0952695) at 4°C and processed within 24 hours of isolation. Histologically normal human adult distal lung tissue was obtained from de-identified specimens through the Michigan Medicine Thoracic Surgery Laboratory, kept at 4°C immediately upon isolation, and processed within 24 hours of isolation.

#### Lung ALI explant Cultures

Human fetal lung within the canalicular stage of lung development was utilized for lung ALI explant cultures (specifically 15 – 18.5 weeks post conception). Small pieces (< 0.5 cm diameter) were dissected from distal regions of the lung and placed on Nucleopore Track-Etched Membrane disks (13 mm, 8 µm pore, polycarbonate) (Sigma, Cat#WHA110414) floating on top of 500 µl of human lung ALI explant media (Advanced DMEM/F-12 (Thermo Fisher, Cat#12634010), 2 mM Glutamax (Thermo Fisher, Cat#35050061), 15 mM HEPES (Corning, Cat#25060CI), 1X B27 Supplement (Thermo Fisher, Cat#17504044) 1X N-2 Supplement (Thermo Fisher, Cat#17502048), 100U/mL penicillin-streptomycin (Thermo Fisher, Cat#15140122)) in a 24 well tissue culture plate (Thermo Fisher, Cat#12565163). Where indicated, 1 µM A-8301 (APExBIO Cat#A3133), 100 ng/mL rhTGFβ1 (R&D Systems Cat#240-B-002), 100 ng/mL rhNOGGIN (produced in-house) or 100 ng/mL BMP4 (R&D Systems Cat#314-BP-050) was added to human lung ALI explant media.

#### BTP-organoid establishment and maintenance

Primary BTP-organoid cultures from 15 – 18.5 wks post conception lung tissue were established and maintained as previously reported^3, 21, 23^. BTP-organoids were maintained in maintenance media (described below) under 8 mg/mL Matrigel (Corning Cat#354234), fed every three days and passaged 1:3 every 7 – 10 days by needle sheering.

#### Needle sheering

Organoids are needle sheered in preparation for passaging by passing the culture through a 27-gauge needle 3 times in 1 mL of progenitor maintenance media (described below) resulting in the fragmentation of organoids.

#### Differentiation of BTP-organoids to AT2-like organoids

Differentiations were performed on BTP-organoids at passage three by removing maintenance media consisting of: DMEM/F-12 (Corning, Cat#10-092-CV), 100U/mL penicillin-streptomycin (Thermo Fisher, Cat#15140122), 1X B-27 supplement (Thermo Fisher, Cat#17504044), 1X N2 supplement (Thermo Fisher, Cat#17502048), 0.05% BSA (Sigma, Cat#A9647), 50µg/mL L-ascorbic acid (Sigma,Cat#A4544), 0.4 µM 1- Thioglycerol (Sigma, Cat#M1753), 50nM all-trans retinoic acid (Sigma, Cat#R2625), 10ng/mL recombinant human FGF7 (R&D Systems, Cat#251-KG), and 3µM CHIR99021 (APExBIO, Cat#A3011) and replacing with differentiation media. For data in figure 2 multiple differentiation medias were tested, all of which consisted of maintenance media with the addition of 1 µM A-8301 and/or the addition of 100 ng/mL BMP4 and/or removal of all trans retinoic acid. B-27 supplement without vitamin A (Thermo Fisher, Cat# 12587010) was used instead of full B-27 supplement in conditions in which all trans retinoic acid was removed. For data in all Figure 3 – 6 differentiation media consisted of common components: DMEM/F12, 100U/mL penicillin-streptomycin, 1X B-27 supplement without vitamin A, 1X N2 supplement, 0.05% BSA, 50µg/mL L-ascorbic acid, 0.4 µM 1-Thioglycerol. 10ng/mL recombinant human FGF7, and 3µM CHIR99021. To make CK + AB differentiation media 1 µM A-8301 and 100 ng/mL BMP4 was added. To make CKDCI differentiation media 50 nM Dexamethasone (Sigma, Cat#D4902), 100 nM 3-isobutyl-1-methylxanthine (Sigma, Cat#I5879) and 100 nM 8- Bromoadensoine 3’,5’-cyclic monophosphate sodium salt (Sigma, Cat#B7880) was added. Differentiations were fed every three days and passaged at seven day intervals by needle sheering.

#### Expansion of AT2-like organoids

After 21 days of differentiation in CK + AB or CK + DCI organoids were passaged and fed with SFFF or SFFF without FGF10 consisting of: Advanced DMEM/F12, 2 mM Glutamax, 1X B27 supplement, 100 U/mL penicillin streptomycin, 15 mM HEPES, 0.05% BSA, 10 µM TGFβ inhibitor SB43152 (APExBIO, Cat#A8249), 1 µM p38 MAP kinase inhibitor BIRB796 (APExBIO, Cat#A5639), 3 µM CHIR99021, 50 ng/mL rhEGF (R&D Systems, Cat#236-EG) with (SFFF) or without the addition of 10 ng/mL FGF10 (produced in-house). Organoids were passaged every 7 – 14 days at a ratio of 1:6 by needle sheering^96^.

#### Primary AT2 Organoid Establishment and Maintenance

Distal lung sections from a single patient were minced using a scalpel. Minced lung was enzymatically dissociated to a single cell suspension using 1 mg/mL collagenase A (Roche, Cat# 10103578001), 2-4U/mL elastase (Worthington, Cat# LS002274), and 0.1 mg/mL DNAse (Roche, Cat#10104159001), filtered through a 100 uM cell strainer, subjected to red blood cell lysis (Roche, Cat #11814389001), washed with PBS, and seeded into Matrigel. After two passages, cultures were subjected to FACS (see below: *Fluorescence activated Cell Sorting*). AT2 cells were isolated on the basis of positive HTII-280 staining and reseeded into Matrigel with primary AT2 organoid media consisting of: Advanced DMEM/F12, 2 mM Glutamax, 1X B27 supplement, 100 U/mL penicillin-streptomycin, 15 mM HEPES, 0.05% BSA, 10 µM SB43152 (APExBIO, Cat#A8249), 1 µM BIRB796 (APExBIO), 3 µM CHIR99021, 50 ng/mL rhEGF and 10 ng/mL rhFGF10 (Cite Katsura). Organoids were expanded in primary AT2 organoid media and passaged every 3-4 weeks at a ratio of 1:2-3 by TrypLE-mediated dissociation.

#### Preparation of tissue, explant and organoids for Fluorescence *In Situ* mRNA Staining and Protein Immunofluorescent (IF) Staining

All samples processed were fixed for 24 hours in 10% Neutral Buffered Formalin at room temperature with gentle agitation. Samples were washed 3x with UltraPure DNase/RNase-Free Distilled Water (Thermo Fisher, Cat#10977015) and dehydrated through an alcohol series consisting of 25%, 50%, 75% and 100% methanol, followed by 100% and 70% ethanol. For tissue and explants each step was performed for at least 1 hour. For organoids each step was performed for at least 15 minutes. In the case of explants and organoids, specimens were embedded in Histogel (VWR Cat# 83009-992) prior to paraffin processing. Samples were paraffin processed in an automated tissue processor through the following series: 70%, 80%, 2x 95%, 3x 100% ethanol, 3x xylene and 3x paraffin with 1 hour for each step. Tissue was embedded into paraffin blocks and cut into 5 µm-thick sections onto charged glass slides using a microtome. Slides were baked for 1 hour at 60 °C immediately prior to staining.

#### Fluorescence *In Situ* mRNA Hybridization (FISH)

FISH was performed using the RNAscope Multiplex Fluorescent V2 assay (ACDBio, Cat# 323100) using TSA- Cy3 (Akoya Biosciences Cat#NEL744001KT) and TSA-Cy5 (Akoya Biosciences, Cat# NEL745E001KT) according to the manufacturer’s recommendations. Protease treatment and Antigen retrieval were performed for 6 and 15 minutes respectively. For Protein Immunofluorescent co-stains, slides were washed in 1x Phosphate Buffered Saline (PBS) (Corning Cat#21-040) after final HRP-Blocker treatment and washes and immediately put into blocking solution for 1 hour, followed by primary and secondary antibody stains as described in the Protein IF Staining protocol below.

#### FISH Quantification

FISH foci were quantified using a custom automated image analysis pipeline in NIS-Elements AR v5 (Nikon).

Nuclei were first segmented and cell borders were estimate by the ‘GrowObjects’ function. Thresholding was then performed to identify RNA foci and the number of foci in each cell was recorded. The lumen of epithelial cells was labelled manually and cells were automatically identified as epithelial based on proximity to lumen. SOX9 immunofluorescent signal for each nuclei was thresholded to distinguish SOX9-positive bud tip and SOX9-negative stalk cells. For each mesenchymal cell the distance (center to center) of the closest SOX9 positive and SOX9-negative epithelial cell was recorded. Distance values were used to categorize if a mesenchymal cell was nearest a SOX9-positive cell, a SOX9-negative cell, or far away (>50 µm) from both. Quantification was performed on 3x field of views per timepoint at 40x magnification.

#### Protein IF Staining

Slides were treated with 2x HistoClear II (National Diagnostics, Cat#HS-202) washes, then rehydrated through washes in 100%, 95%, 70%, 30% ethanol for 4 minutes each, with buffer exchanges performed halfway through washing. Then, slides were washed 2x 5 minutes with ddH20. Antigen retrieval in 1X Sodium Citrate Buffer (100mM trisodium citrate (Sigma, Cat#S1804), 0.5% Tween-20 (Thermo Fisher, Cat#BP337), pH 6.0) for 20 minutes at 99 °C. After washing 3x in ddH20 slides were blocked for 1 hour with blocking solution: 5% normal donkey serum (Sigma, Cat#D9663), 0.1% Tween-20 in PBS. Slides were then incubated in primary antibodies diluted in blocking solution in a humidified chamber at 4°C overnight. Slides were washed 3x in 1X PBS for 10 minutes each. Slides were incubated with secondary antibodies and DAPI (1µg/mL) diluted in blocking solution for 1 hour, then were washed 3x in 1X PBS for 5 minutes each. Slides were mounted in ProLong Gold (Thermo Fisher, Cat#P369300) and imaged within 2 weeks. Primary and secondary antibodies used in this study are available in Supplementary Table 1.

#### Whole Mount Protein IF Staining

Organoids were fixed in 10% NBF overnight at room temperature on a rocker. Tissue was then washed three times for 2 hours in Organoid Wash Buffer (OWB) (0.1% Triton X-100, 0.2% BSA, 1x PBS) at RT on a rocker. Organoids were then submerged in CUBIC-L (TCI Chemicals Cat#T3740) for 48 hours at 37°C. Organoids were then permeabilized with permeabilization solution (5% Normal Donkey Serum, 0.5% Triton X-100, 1x PBS) for 24 hours at 4°C. Organoids were washed 1x with OWB and then incubated with primary antibody (diluted in OWB) for 24 hours at 4°C. Organoids were then washed 3x with OWB and secondary antibody (diluted in OWB) was added for 2 hours at RT. Organoids were washed an additional 3x with OWB and then cleared in CUBIC-R (TCI Chemicals Cat#T3741) with 1 µg/mL DAPI. Cleared organoids were mounted on slides with Secure-Seal Spacers (Invitrogen Cat#S24737) to accommodate 3-dimensional imaging.

#### Preparation and transmission electron microscopy of organoids

Transmission electron microscopy sample preparation was performed by the University of Michigan BRCF Microscopy and Image Analysis Laboratory. Samples were fixed in 3% glutaraldehyde/3% paraformaldehyde in 0.1M cacodylate buffer (CB), pH 7.2. Samples were washed 3 times for 15 minutes in 0.1M CB and then kept for 1 hour on ice in 1.5% K_4_Fe(CN)_6_ + 2% OsO_4_ in 0.1M CB. Samples were washed 3x in 0.1M CB, followed by 3x in 0.1M Na_2_ + Acetate Buffer, pH 5. Staining contrast enhancements by 1 hour treatment with 2% Uranyl Acetate + 0.1M Na_2_ + Acetate Buffer, pH 5.2. Samples were then processed overnight in an automated tissue processor, including dehydration from H_2_O through 30%, 50%, 70%, 80%, 90%, 95%, 100% ethanol, followed by 100% acetone. Samples were infiltrated with Spurr’s resin at an acetone: Spurr’s resin ration of 2:1 for 1 hour, 1:1 for 2 hours, 1:2 for 16 hours, and absolute Spurr’s resin for 24 hours. After embedding and polymerization, samples were sectioned on an ultramicrotome. TEM samples were imaged on a JEOL JEM 1400 PLUS TE microscope.

#### RNA extraction, Reverse Transcription and RT-qPCR

Organoids were dislodged from Matrigel using a P1000 tip, pelleted in a 1.5 mL tube and flash frozen by placing the tube in a small amount of liquid nitrogen with minimal residual media. RNA was isolated from frozen pellets using the MagMax-96 Total RNA Isolation kit (Thermo Fisher, Cat#AM1830) according to the manufacturer’s recommendations. RNA quality and yield was determined on a Nanodrop 2000 spectrophotometer. Reverse Transcription was performed in triplicate for each biological replicate using the SuperScript VILO cDNA kit with 200ng RNA per reaction. After Reverse Transcription, cDNA was diluted 1:2 with DNAse/RNAse free water and 1/40^th^ of the final reaction was used for each RT-qPCR measurement. RT-qPCR measurements were performed using a Step One Plus Real-Time PCR System (Thermo Fisher, Cat#43765592R) using QuantiTect SYBR Green qPCR Master Mix (Qiagen, Cat#204145) with primers at a concentration of 500 nM. Sequences for RT-qPCR primers used in this manuscript are in Supplementary Table 2.

#### Fluorescence-activated Cell Sorting (FACS)

Organoids were retrieved from Matrigel by mechanical dissociation with a P1000, washed 2x with 1 mL PBS and, resuspended with TrypLE and incubated at 37°C until a single cell suspension was obtained with light pipetting (typically 10 minutes). Cells were washed 3x with 1 mL of FACS buffer consisting of PBS, 2% BSA and 10 µM Y-27632 (APExBIO, Cat#3008) and then filtered through a 30 µM mesh. Cells were stained for 1 hour on ice in FACS buffer with a 1:60 dilution of anti HTII-280 IgM antibody. Cells were washed 3x with 1 mL of FACS buffer before 30 minutes of staining in FACS buffer with a 1:1000 dilution of anti-mouse IgM Alexaflour-488 secondary antibody (Jackson Immunoresearch, Cat#715545140) on ice. After 3x washes with 1 mL of FACS buffer cells were suspended in FACS buffer with 1:4000 dilution of DAPI. Live cells (determined by exclusion of DAPI) positive for HTII-280 were identified by 488 emission intensity notably higher than primary only, secondary only controls and absence of 405 (DAPI) emission. All steps were carried out at 4°C unless otherwise indicated. Cells were pelleted at 500xG for 5 minutes in swing bucket rotors. FACS was performed on either a Sony MA900 or the ThermoFisher Bigfoot Spectral Cell Sorter.

#### Preparation of Lung ALI explant cultures for single cell RNA-sequencing (scRNA-seq)

For scRNA-sequencing of Lung ALI explant cultures, the outside rind of n = 3 explants at each timepoint (0, 3, 6, 9 and 12 days) was micro dissected and minced using a No.1 Scalpel, discarding the center of the explant, which appeared necrotic in later timepoints. Pooled explants at each timepoint (0, 3, 6, 9 and 12 days) were dissociated using the Neural Tissue Dissociation Kit (P) (Mitenyi Biotec, Cat#130092628). Briefly minced explant tissue was resuspended in Mix 1 and incubated at 37°C for 15 minutes. Mix 2 was then added and incubated for 10 minutes at 37°C. Cells were agitated by P1000 pipetting and then returned to the incubator for additional 10 minute incubations at 37°C, followed by P1000 pipetting until a single cell suspension was achieved (∼30 minutes). Obtained single cell suspension was filtered through a 70 µm Flowmi Cell Stainer (Sigma, Cat#BAH136800070) and then resuspended in Red Blood Cell Lysis Buffer (Roche, 11814389001) for 15 minutes at 4°C. After Red Blood Cell Lysis, cells were washed twice with 2mL 1X HBSS + 1% BSA and then resuspended in Cryostor-CS10 (Sigma, Cat#C2874) for storage in liquid nitrogen. All Lung ALI explant culture samples were thawed and co-submitted for sequencing on the same day. Thawing of cells prior to sequencing consisted of adding 1:1 increments of RPMI + 10% FBS drop-wise, with 1 minute pauses every time the volume doubled, until a total volume of 32 mLs was achieved. Cells were then pelleted and resuspended in 1 mL HBSS + 1% BSA and passed through a 40 µm Flowmi Cell Strainer (Sigma, Cat#BAH136800040), counted on a hemocytometer and submitted at 1000 cells/µl in HBSS + 1% BSA to the University of Michigan Advanced Genomics Core for library preparation by the Chromium Next GEM Single Cell 3’ GEM, Library and Gel Bead Kit v3.1 (10x Genomics, Cat#PN1000128) targeting 7500 cells. scRNA sequencing libraries were sequenced using a NovaSeq 6000 with S4 300 cycle reagents (Illumina, Cat#20028312). Cells were pelleted by spinning at 500xG for 5 minutes in a swing-bucket centrifuge. All steps were carried out using tips coated in HBSS + 1% BSA and pre-chilled (4°C) buffers and equipment.

#### Preparation of Organoids for scRNA-seq

Organoids were dislodged from Matrigel by mechanical dissociation using a p1000 pipette tip. Pelleted organoids were resuspended in TrypLE Express (Thermo Fisher, Cat #17105041) and incubated at 37°C, pipetting gently at 5 minute intervals until a single cell suspension is obtained (∼10 minutes). Cells were washed 3x with Hanks Balanced Salt Solution (HBSS) (Thermo Fisher, Cat #14175095) + 1% BSA and then passed through a 40 µm FlowMi Cell Strainer. Cells were counted on a hemocytometer and then resuspended at 1000 cells/µl in HBSS + 1% BSA for submission to the the University of Michigan Sequencing Core, which prepared libraries using the Chromium Next GEM Single Cell 3’ GEM, Library and Gel Bead Kit v3.1 (10x Genomics, Cat#PN1000128) targeting 3500 cells. scRNA-sequencing libraries were sequenced using a NovaSeq 6000 with S4 300 cycle reagents. Cells/organoids were pelleted by spinning at 500xG for 5 minutes in a swing-bucket centrifuge. All steps were carried out using tips coated in HBSS + 1% BSA and pre-chilled (4°C) buffers and equipment.

#### Expression and purification of recombinant human FGF10

The expression plasmid for recombinant human FGF10 (pET21d-FGF10) was a gift from James A Bassuck (Bagai et al., 2002). FGF10 expression was induced by the addition of isopropyl-1-thio-B-D-galactopyranoside to Rosetta^TM^2(DE3)pLysS carrying pET21d-FGF10 in 2x YT medium (BD Biosciences, Cat#244020) with Carbencillin (50 µg/mL) and Chloramphenicol (17 µg/mL). FGF10 was purified using a HiTrap-Heparin HP column (GE Healthcare, Cat#17040601) with step gradients of 0.2 to 0.92 M NaCl. Purity of FGF10 was assessed by SDS-PAGE gel and activity based on the efficiency to phosphorylate ERK1/2 in A549 cells (ATCC, Cat#CCL-185)

#### scRNA-seq Analysis

##### Overview

To identity clusters of cells with similar gene expression within scRNA-sequencing datasets we processed CellRanger filtered matrices using Seurat v4.0^49^ in RStudio v1.4 running R v4.2. The general workflow involves filtering for high quality cells, normalizing counts to read depth, log transformation and scaling the normalized count data, identification of variable genes between cells, identification of principal components, batch correction (if applicable), uniform manifold approximation and Louvain clustering of cells.

##### Quality Control

Cells were filtered for under or over (likely doublets) complexity by filtering based on number of features detected and for cell viability/quality based on the percentage of mitochondrial reads in a cell’s transcriptome. Cells not conforming to the following parameters were removed from analysis: Fig. 1a,b,c, Supplementary Fig. 1a,b – features detected : >500, < 5000, mitochondrial reads: <10%; Fig. 1j,k,l and Supplementary Fig.1f,g,h,I,j,k,l – features detected: >1000, <12000,;mitochondrial reads: <10%, Fig. 3 – features detected: >2500, <10000; mitochondrial reads: < 20%, Fig. 5 – features detected: >2000, <10000; mitochondrial reads: < 20%; Fig. 6 – features detected: >2500, <10000; mitochondrial reads; < 20%.

##### Gene Expression Visualization and Differential Expression

Prior to visualization or analysis gene expression counts were normalized to total counts for each cell, multiplied by factor of 10000 and natural log transformed. Significance of gene expression differences was determined by Wilcoxon Ranked Sum test and limited to genes with at least 25% of cells expressing within at least one group compared, and log 2 transformed normalized count differences greater than 0.25.

##### Batch Correction

For the analysis of ALI explant cultures in Fig. 1 and Supplementary Fig. 1 samples from multiple days of culture were batch corrected using the Seurat implementation of reciprocal PCA analysis. For the analysis of CK + AB treated (Fig. 3) and CK + DCI treated (Fig. 6) BTP organoids, day 0 BTP organoid samples were integrated with day 1, day 6 and day 21 treated samples using FastMNN^97^ through the SeuratWrappers package. For Fig. 5i,j AT2-mapping cells (part f) from primary AT2 organoids, CK + AB AT2-like organoids and CK + DCI AT2-like organoids were also batch corrected using FastMNN. In all cases batch correction was performed using the top 30 most variable principal components.

##### Dimensional Reduction and Clustering

For samples processed individually, SCTransform was used for normalization and scaling prior to identification of variable features and reduction by Principal Component analysis (PCA). For batch corrected samples the corrected gene expression matrix was scaled and then used as input for PCA reduction. Using the top principal components (PCs) a neighborhood graph was constructed from 20 nearest neighbors focusing on highly variable PCs (Supplementary Fig. 1f,g,h,l - 18 PCs; Fig. 1j,k,l and Supplementary Fig. 1i,j,k - 7 PCs; Supplementary Fig. 3e,f,g - 18 PCs; Fig. 3 - 30 PCs; Fig. 5a,b and Supplementary Fig. 5a,b,c,d - 16 PCs; Fig. 5i,j - 30 PCs; Supplementary Fig. 6d,e,f - 16 PCs; Fig. 6b,c,d,e,f and Supplementary Fig. 6g - 30 PCs). Clusters were identified using the Louvain algorithm with specified resolution (Supplementary Fig. 1f,g,h,i – 0.5; Fig. 1j,k,I and Supplementary Fig. 1i,j,k – 0.4; Supplementary Fig. 3e,f,g– 0.5; Fig. 3 – 0.3; Fig. 5a,b and Supplementary Fig. 5a,b,c,d – 0.5; Fig. 5i,j – 0.2; Supplementary Fig. 6d,e,f – 0.5; Fig. 6b,c,d,e,f and Supplementary Fig. 6g – 0.3).

##### Trajectory Analysis

Batch corrected scRNA-seq timecourse data from CK + AB and CK + DCI treated BTP organoids were clustered at increased resolution prior to trajectory analysis (Fig. 3k – 1.0; Fig. 6e – 1.0). Trajectory analysis was performed using Slingshot^44^ with the start cluster determined by the cluster with the highest proportion of cells from BTP organoids. Trajectories terminating in areas of high AT2 marker gene expression at day 21 were chosen as the trajectory corresponding to BTP organoid cells differentiating towards AT2-like identity.

##### Gene Set Module Scoring

Gene set module scoring was performed using the Seurat v4 implementation of the gene set method developed by Tirosh et al.,^45^. Briefly control genes (100 control genes for each module gene) are randomly selected from a bin of similar expressed genes and then expression levels of genes in the module set relative to control genes are calculated. To define an AT2 differentiation gene module in Fig. 3m, scRNA-sequencing data from primary AT2 organoids in SFFF without FGF10 media was merged with scRNA-sequencing data from BTP organoids. Low resolution (0.1) Louvain clustering identified BTP organoids and primary AT2 organoids as distinct clusters of cells. Differential expression analysis was performed between each cluster to identify the top 199 genes significantly enriched in primary AT2 organoids over BTP organoids (Supplementary Table 3). To define a gene module for *in vivo* AT2 cells in Fig. 5L-M, 205 markers reported to be AT2 enriched relative to all other lung cell populations in scRNA-sequencing data from adult lungs were used^31^. 6 of these genes were not detected in our sequencing data, leaving 199 in the scoring module (Supplementary Table 4).

##### Reference-based Mapping

Extracted epithelial cells from scRNA-sequencing of human proximal and distal airways were downloaded from Gene Expression Omnibus (GSE178360)^48^ and used as reference. Data was normalized and variable features were identified in organoid data, and pre-existing variable features in the reference were used. Reference PCA was projected onto query data using the 20 most variable PCs to identity anchors which were applied for cell identity assignment in the query and additionally for projecting query data on the reference UMAP utilizing the MapQuery function in Seurat v4.

### Statistical Analysis

Statistical analysis was performed in PRISM 9 (GraphPad Software).

### FACS

For Fig. 4g statistical significance (p) between the percent of HTII-280-positive cells in SFFF with and without FGF10 was calculated by paired Student’s t-test.

### Image Quantification

For Fig. 1e and f statistical significance (p) was calculated between the # of foci per cell within the same cell population at different timepoints or between different populations at the same timepoint by one-way ANOVA with Bonferroni correction.

### RT-qPCR

For RT-qPCR data arbitrary units (AUs) of gene expression was first calculated using the following equation: 2^(*GAPDH*Ct − TargetCt)^ X 10,000. For Supplementary Fig. 2b, Supplementary Fig. 3d, Fig. 4i, Supplementary Fig. 5e and Fig. 6i the AUs for each biological replicate were calculated from the mean of technical replicates and a ratio paired t-test was performed between the two conditions compared to determine statistical significance (p). For Fig. 2e AUs were normalized to AUs of expression in progenitor media for each biological replicate to

obtain fold change. Repeated measures one-way ANOVA was performed with Dunnett’s correction on linearized (log-transformed) fold change values comparing all experimental conditions to progenitor medium to determine p.

## Data Availability

### Lead Contact

Requests for materials used in this study should be directed to Jason R. Spence at.

## Materials availability

This study did not generate any tools or reagents.

## Data and code availability

Single Cell Sequencing data used in this study is available at EMBL-EBI ArrayExpress, Gene Expression Omnibus or Synapse.org. **EMBL-EBI ArrayExpress** Single-cell RNA sequencing of human fetal lung (<u>E-MTAB-8221</u>)^23^, human cananicular stage lung ALI explants (E-MTAB-12959) (this study), and human lung organoids (E-MTAB-12960) (this study). **Gene Expression Omnibus**: Single-cell RNA sequencing of micro dissected human distal airways (<u>GSE178360</u>)^48^, **Synapse.org**: Human Lung Cell Atlas (<u>syn21041850</u>)^31^.

Code for analysis of scRNA-sequencing data is available at: https://github.com/jason-spence-lab/Frum-et-al.

## Acknowledgements

This study was made possible by funding from the following sources: CZF2019-002440 from the Chan Zuckerberg Initiative DAF, an advised fund from the Silicon Valley Community Foundation and by NIH-NHLBI R01HL119215 and NHLBI R01HL166139 to J.R.S.. T.F. was additionally supported by NIH NIDCR T32DE007057. P.P.H is additionally supported by the Rogel Cancer Center Fellowship. R.F.C.H. is additionally supported by NIH-NHLBI F31HL152531. A.C.S. was supported by 5T32GM007863 and is currently supported by NIH-NHLBI F30HL156474. I.A.G is supported by NIH-NICHD 5R24HD000836. J.Z.S. is supported by NIH-NIDDK R01DK120623.

We thank the University of Michigan Advanced Genomics core, the University of Michigan Microscopy core and the University of Michigan Flow Cytometry core for assistance with experiments in this manuscript. We also acknowledge the University of Washington Laboratory of Developmental Biology for access to lung tissue. Additionally, we acknowledge current members of the Spence Laboratory for helpful insight and discussion during this study and Lindy K. Brastrom for critical reading of the manuscript.

## Author Contributions

T.F. and J.R.S. conceived the study. J.R.S. supervised the research. T.F. designed, performed and interpreted experiments, computational analysis and assembled figures. P.P.H. established primary AT2 organoid lines and assisted with experiments. I.A.G and J.Z.S. contributed critical resources to the project. C.J.Z, R.C.F.H, A.C.S., A.A., O.U., S.G.C. and Y.Z. assisted with experiments. T.F. and J.R.S. wrote the manuscript. All authors read, contributed feedback, and approved the manuscript.

---

**Supplementary Fig. 1:**
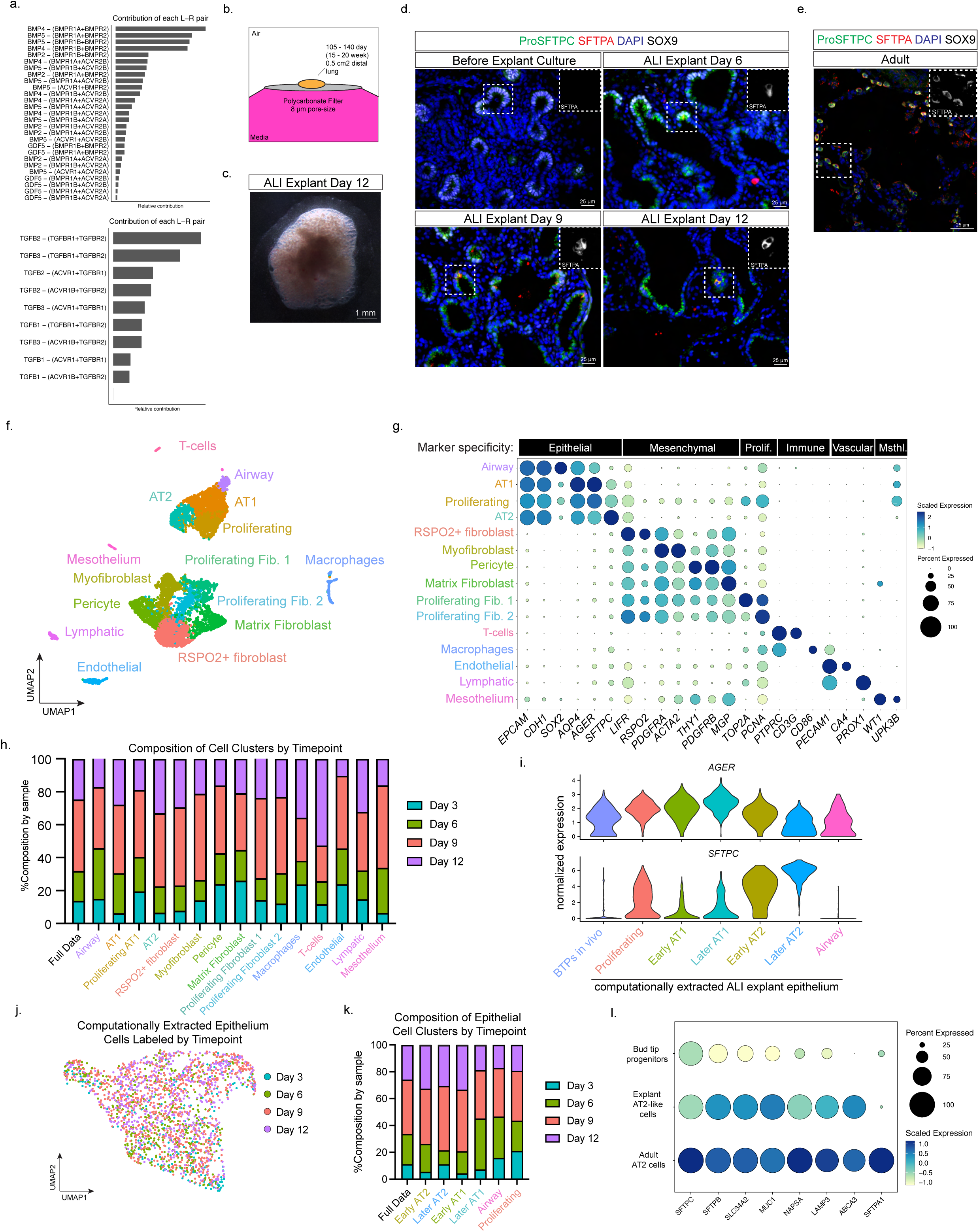
Ligand-receptor pairs contributing to BMP- and TGFβ-signaling during fetal lung development and characterization of canalicular stage lung air-liquid interface (ALI) explant culture. Related to Figure 1. a) Ligand receptor pairs contributing to cell-cell signaling predictions in Figure 1a. b) Schematic of lung explant air-liquid interface culture. 0.5 cm^2^ pieces of distal canalicular stage lung are cultured on polycarbonate filters that float on growth-factor and serum-free media. c) Bright-field image of explant ALI cultures at day 12. Media is observed around the base of the explant, but otherwise the explant is exposed to air. d) Immunofluorescent staining images of AT2 markers (ProSFTPC, SFTPA) and BTP marker SOX9 in canalicular stage lung explants before and after 6, 9 or 12 days of explant ALI culture. e) Immunofluorescent staining images of AT2 markers (ProSFTPC, SFTPA) and BTP marker SOX9 in adult alveoli. Adult AT2s uniformly co-express ProSFTPC and SFTPA and do not stain positive for nuclear SOX9. f) UMAP visualization of Louvain clustering of all cells from day 3, day 6, day 9 and day 12 explants. Cluster identities were assigned on the basis of marker expression in part g. g) Dot plot showing cluster specific marker expression. Marker specificity on the top row denotes cell type/state indicated by unique expression of markers shown. Prolif. = proliferation. Msthl. = mesothelium. h) Quantification of the percent contribution of each timepoint to clusters identified in integrated scRNA-sequencing data from ALI explant culture. The contribution of each timepoint to the full dataset is shown in the leftmost column. i) Comparison of *AGER* (top) and *SFTPC* (bottom) expression between BTPs in lung tissue prior to ALI explant culture, and clusters identified in computationally extracted epithelial cells. j) UMAP of computationally extracted epithelial cells from scRNA-sequencing of ALI explant culture with cells colored by days of ALI explant culture. k) Quantification of the percent contribution of each timepoint to clusters identified in integrated computationally extracted epithelium from ALI explant scRNA-sequencing data. The contribution of each timepoint to the full dataset is shown in the leftmost column. l) Dot plot comparing expression of AT2 markers in BTPs, explant AT2-like cells and primary AT2 cells. Explant AT2-like cells express higher AT2 markers than bud tip progenitors and less than adult AT2 cells.

**Supplementary Fig. 2:**
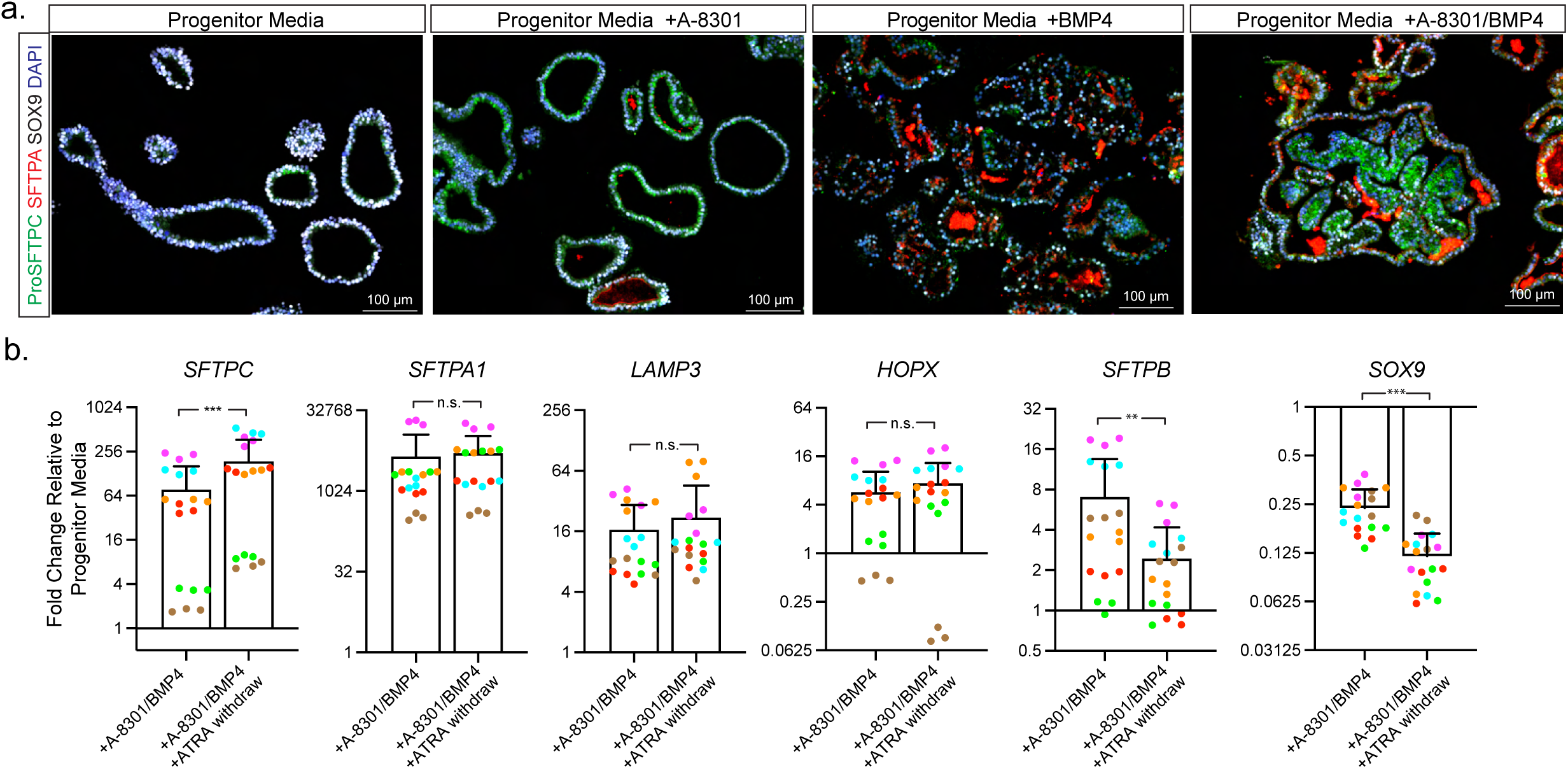
AT2 and BTP marker expression in BTP organoids under TGFβ-inhibition and BMP-activation and the effect of all-trans retinoic acid (ATRA) on AT2 differentiation of BTP organoids. Related to Figure 2. a) Immunofluorescent staining images of AT2 markers (ProSFTPC and SFTPA) and BTP marker SOX9 in BTP organoids cultured in progenitor media or progenitor media with addition of TGFβ inhibitor A-8301 and BMP activator BMP4 alone or simultaneously for seven days. b) RT-qPCR measurements comparing AT2 markers (*SFTPC, SFTPA1, LAMP3, HOPX*, *SFTPB)* or BTP marker *SOX9* in response to simultaneous TGFβ-inhibition and BMP-activation in the presence (+A-8301/BMP4) or absence (+A8301/BMP4 +ATRA Withdraw) of ATRA for seven days. Values shown are fold change relative to BTP organoids maintained in Progenitor media for each technical replicate color-coded by biological replicate. Statistical comparison (p) was computed by ratio paired t-test on the mean arbitrary units of expression for each biological replicate calculated from the mean of technical replicates. (* = p < 0.05, ** = p < 0.01, *** = p < 0.001, n.s. = p > 0.05.). Error bars = standard deviation.

**Supplementary Fig. 3:**
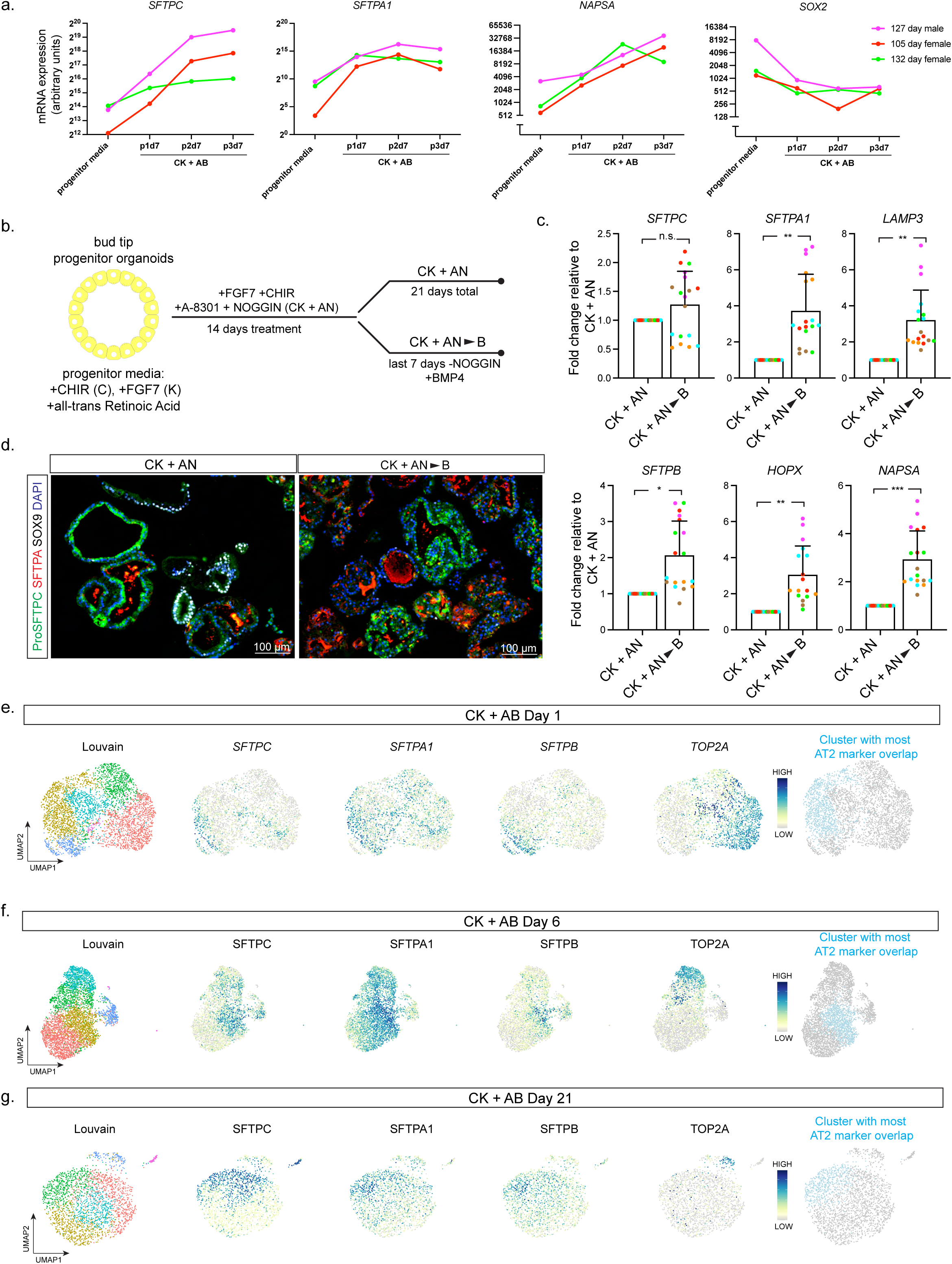
Reproducibility of CK + AB response in multiple BTP organoid lines, evidence for maximal differentiation in the presence of BMP-activation and identification of clusters with the most AT2 marker overlap at each CK + AB treatment timepoint. Related to Figure 3. a) RT-qPCR measurements showing arbitrary units of mRNA expression for AT2 markers (*SFTPC, SFTPA1, NAPSA*) and airway marker *SOX2* in response to CK + AB over the course of 21 days for three BTP organoid lines. b) Schematic of BTP organoid differentiation experiment analyzed by RT-qPCR in part c and Immunofluorescent staining in part d. Modified CK + AB media made to inhibit BMP-signaling rather than activate it by replacing BMP4 with NOGGIN (CK + AN) was applied to BTP organoids for 14 days. Cultures were then divided with half receiving CK + AB and the other half maintained in CK + AN with analysis performed after an additional 7 days (21 days total). c) RT-qPCR measurements of AT2 marker expression in BTP organoids treated as schematized in part b. Values shown are fold change relative to the CK + AN condition for each technical replicate color-coded by biological replicate. Statistical comparison (p) was computed by ratio paired t-test on the mean arbitrary units of expression for each biological replicate calculated from the mean of technical replicates. (* = p < 0 .05, ** = p < 0.01, *** = p < 0.001, n.s. = p > 0.05). Error bars = standard deviation. d) Immunofluorescent staining for AT2 markers (ProSFTPC, SFTPA) and BTP marker SOX9 in BTP organoids differentiated under the conditions schematized in part b. e) UMAP visualization of Louvain clustering and gene expression for AT2 markers (*SFTPC, SFTPA1, SFTPB)* and proliferation marker *TOP2A* in CK + AB treated BTP organoids after 1 day. The cluster with the highest overlapping expression of *SFTPC*, *SFTPA* and *SFTPB* is highlighted in the rightmost plot. f) UMAP visualization of Louvain clustering and gene expression for AT2 markers (*SFTPC, SFTPA1, SFTPB)* and proliferation marker *TOP2A* in CK + AB treated BTP organoids after 6 days. The cluster with the highest overlapping expression of *SFTPC*, *SFTPA* and *SFTPB* is highlighted in the rightmost plot. g) UMAP visualization of Louvain clustering and gene expression for AT2 markers (*SFTPC, SFTPA1, SFTPB)* and proliferation marker *TOP2A* in CK + AB treated BTP organoids after 21 days. The cluster with the highest overlapping expression of *SFTPC*, *SFTPA* and *SFTPB* is highlighted in the rightmost plot.

**Supplementary Fig. 4:**
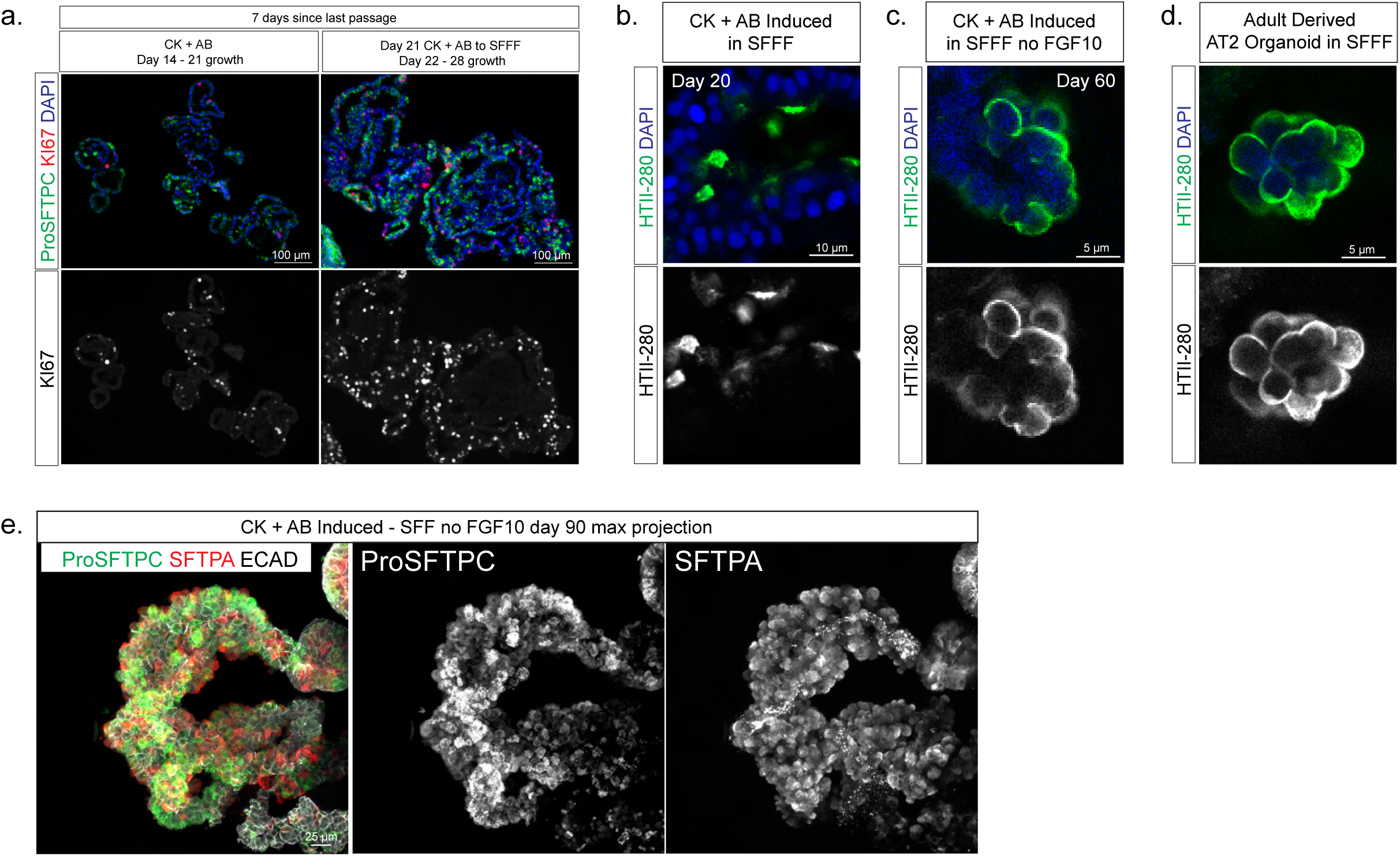
Proliferative and morphological features of CK + AB induced AT2-like organoids in CK + AB, SFFF and SFFF without FGF10 medias. Related to Fig. 4. a) Immunofluorescent staining images of AT2 marker ProSFTPC and proliferation marker KI67 in CK + AB induced AT2-like organoids at the end of the final 7 days of 21 day CK + AB differentiation, or after the first 7 days of switching the culture to SFFF. b,c,d) Immunofluorescent staining images of AT2 marker HTII-280 in CK + AB induced organoids expanded in SFFF for 20 days (b), CK + AB induced organoids expanded in SFFF without FGF10 for 60 days (c) or primary AT2 organoids in SFFF (d). e) Maximum signal intensity projection of confocal images from whole mount immunofluorescent staining of AT2 markers ProSFTPC and SFTPA in CK + AB induced organoids expanded in SFFF without FGF10 for 90 days. ECAD staining is included in the merged image to identify cell boundaries.

**Supplementary Fig. 5:**
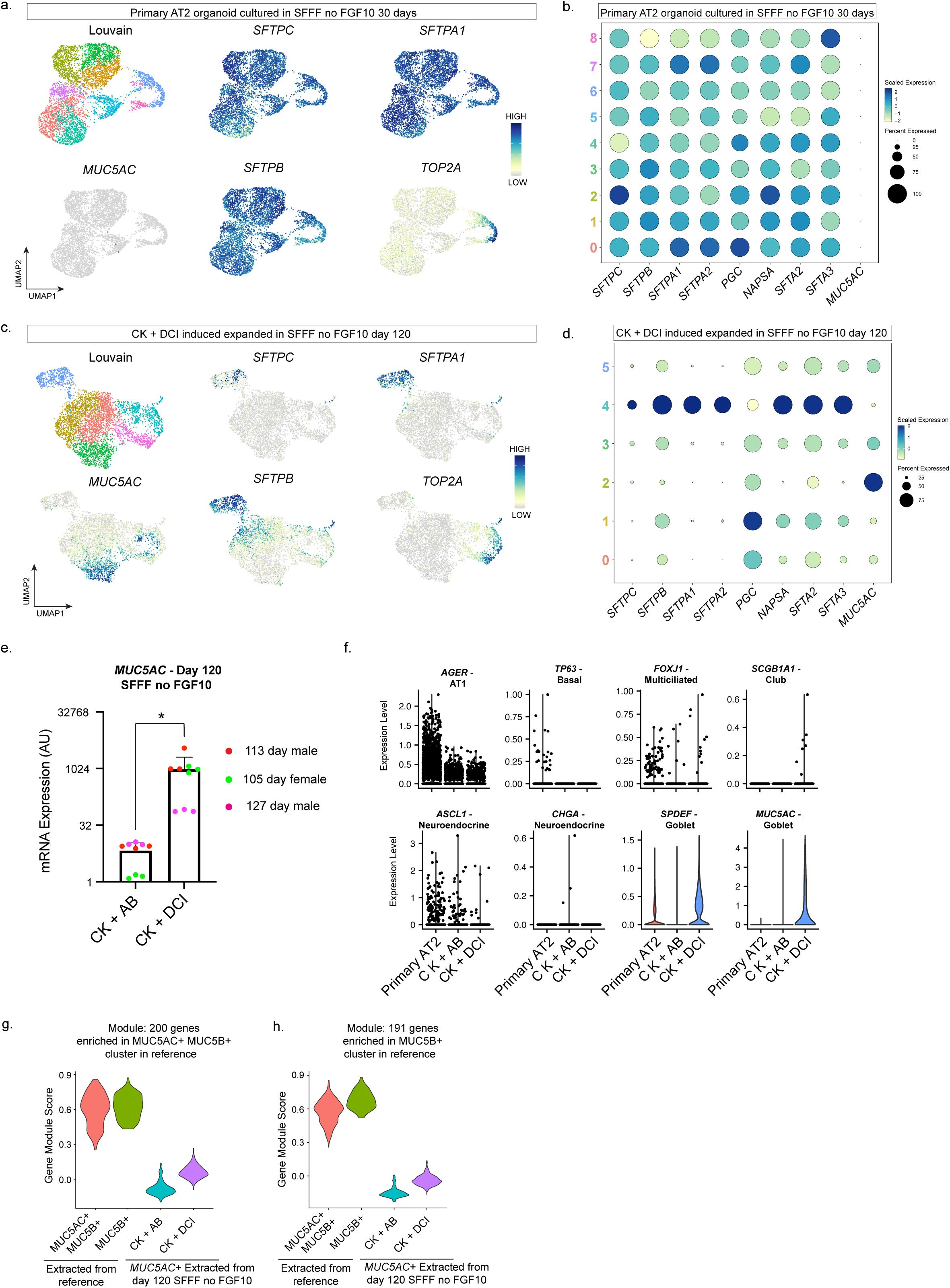
Composition of primary AT2 and CK + DCI differentiated organoids expanded in primary AT2 media for 120 days and RT-qPCR validation of differences between expanded CK + AB and CK + DCI differentiated organoids. Related to Fig. 5. a) UMAP visualization of Louvain clustering and gene expression in primary AT2 organoid cultures established in SFFF for multiple passages and switched to SFFF without FGF10 for 30 days before analysis. AT2 markers (*SFTPC, SFTPA1, SFTPA*), proliferation marker *TOP2A* and goblet cell marker *MUC5AC* are shown. b) Dot plot showing homogenous expression of an expanded panel of AT2 markers and absence of goblet cell marker MUC5AC across Louvain clusters in primary AT2 organoids established in SFFF for several passages and cultured SFFF without FGF10 for 30 days before analysis. c) UMAP visualization of Louvain clustering and gene expression in CK + DCI differentiated organoids expanded for 120 days in SFFF without FGF10. AT2 markers (*SFTPC, SFTPA1, SFTPB*), proliferation marker *TOP2A* and goblet cell marker *MUC5AC* are shown. d) Dot plot showing expression of an expanded panel of AT2 markers and goblet cell marker *MUC5AC* in Louvain clusters in CK + DCI differentiated organoids expanded for 120 days in SFF without FGF10. e) RT-qPCR measurements of *MUC5AC* expression in three BTP organoid lines differentiated with CK + AB or CK + DCI and expanded for 120 days in SFFF without FGF10. Arbitrary units (AU) of gene expression are shown. Statistical significance (p) was calculated by ratio paired t-test on the mean AUs for each biological replicate calculated from three technical replicates. (* = p < 0.05). (* = p < 0 .05, ** = p < 0.01, *** = p < 0.001, n.s. = p > 0.05). Error bars = standard deviation. f) Violin plot comparing expression of markers of non-AT2 lung epithelial cell types in primary AT2 organoids. Relative to goblet cell markers (*SPDEF, MUC5AC*) expression of other non-AT2 lung epithelial cell type markers is not detected (*TP63*), infrequent (*SCGB1A1*, *CHGA*) or less than primary AT2 organoids (*AGER*, *FOXJ1*, *ASCL1*). g) Violin plots showing gene module scores for primary goblet cells extracted from an in vivo reference data set or goblet-like MUC5AC positive cells extracted from day 120 CK +AB and CK + DCI AT2-like organoids in SFFF without FGF10. The gene module was comprised of the top 200 genes enriched in the cluster annotated ‘MUC5AC+ MUC5B+’ in the in vivo reference dataset. h) Violin plots showing gene module scores for primary goblet cells extracted from an in vivo reference data set or goblet-like MUC5AC positive cells extracted from day 120 CK +AB and CK + DCI AT2-like organoids in SFFF without FGF10. The gene module was comprised of all 191 genes enriched in the cluster annotated ‘MUC5B+’ in the in vivo reference dataset.

**Supplementary Fig. 6:**
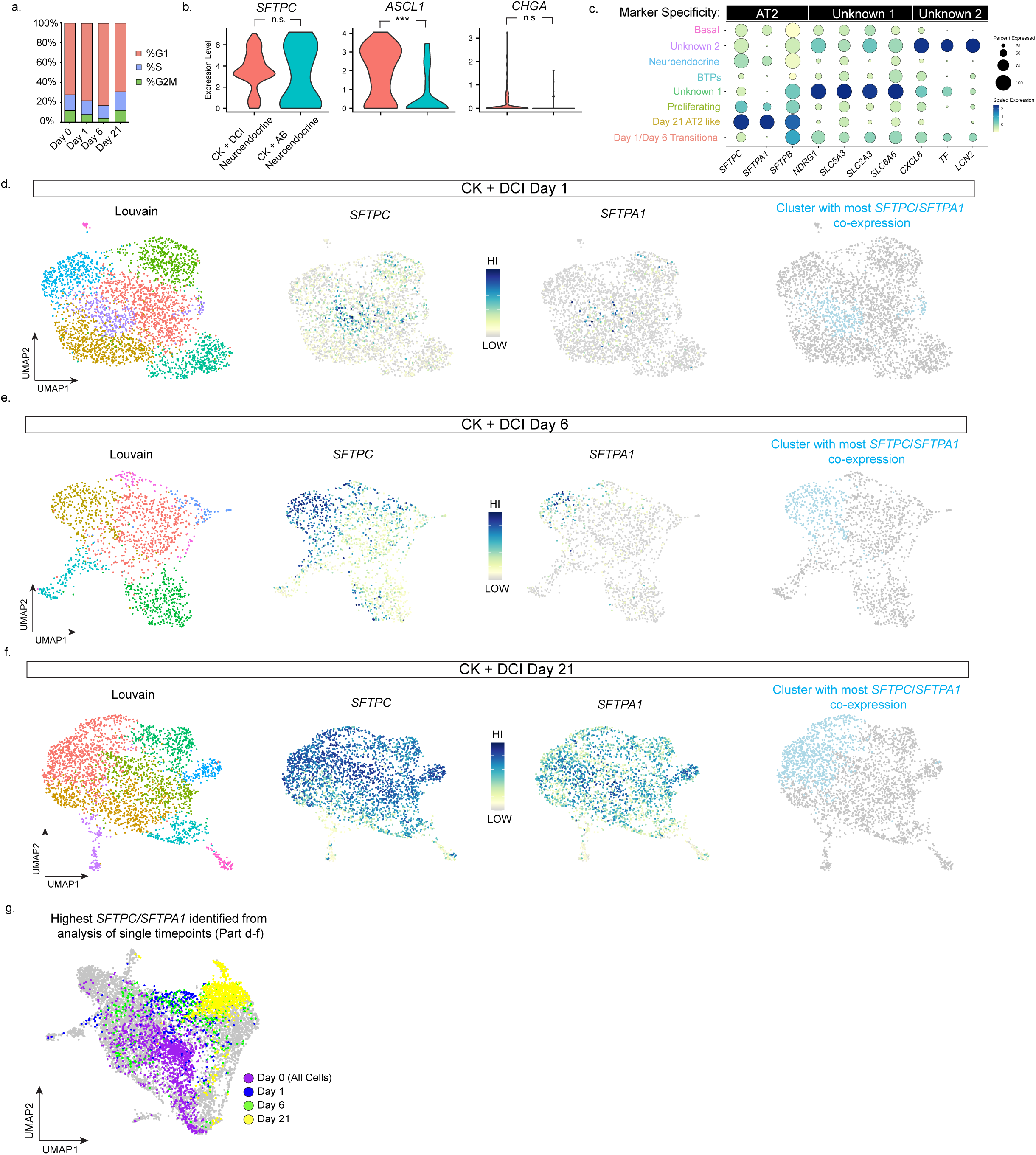
Characterization of the transcription response of BTP organoids to CK + DCI and identification of clusters with the most AT2 marker overlap at each timepoint of CK + DCI treatment. Related to Figure 6. a) Comparison of the percentage of cells in each phase of the cell cycle in BTP organoids (day 0) and at days 1, 6 and 21 of CK + DCI treatment. b) Violin plots comparing AT2 marker *SFTPC* or neuroendocrine marker expression (*ASCL1, CHGA*) in neuroendocrine-like cells present in either CK + DCI or CK + AB treated BTP organoids. Significance of differential enrichment (p) between the ASCL1+ clusters from CK + DCI and CK + AB differentiations was determined by Wilcoxon Rank Sum test (*** = p < 0.001, n.s. = p > 0.05). c) Dot plot showing markers enriched in clusters of unknown identity from integrated scRNA-seq data of CK + DCI treatment time course. d) UMAP visualization of Louvain clustering and gene expression for AT2 markers (*SFTPC, SFTPA1*) in CK + DCI treated BTP organoids after 1 day. The cluster with the highest overlap of shown AT2 markers is highlighted in the rightmost plot. e) UMAP visualization of Louvain clustering and gene expression for AT2 markers (*SFTPC, SFTPA1*) in CK + DCI treated BTOs after 6 days. The cluster with the highest overlapping expression of *SFTPC* and *SFTPA1* is highlighted in the rightmost plot. f) UMAP visualization of Louvain clustering and gene expression for AT2 markers (*SFTPC, SFTPA1*) in CK + DCI treated BTOs after 21 days. The cluster with the highest overlapping expression of *SFTPC* and *SFTPA1* is highlighted in the rightmost plot. g) UMAP visualization of integrated scRNA-seq data from BTP organoids (day 0) and day 1, 6 and 21 of CK + DCI treatment with highest *SFTPC/SFTPA1* co-expressing cells from independent analysis of each scRNA-seq timepoint (part d-f) highlighted and color coded by timepoint.

Supplementary Table 1: Primary and secondary antibodies used for immunofluorescent staining.

| <u>Name</u> | <u>Company</u> | <u>Catalog #</u> | <u>Dilution</u> | <u>Antibody Registry ID</u> |
| --- | --- | --- | --- | --- |
| Primary Antibodies |  |  |  |  |
| Goat anti-SOX9 | R&D Systems | AF3075 | 1:500 | AB_2194160 |
| Rabbit anti-SOX9 | Millipore | AB5545 | 1:500 | AB_2239761 |
| Goat anti-CDH1 | R&D Systems | AF748 | 1:500 | AB_355568 |
| Rabbit anti-ProSFTPC | Seven Hills Bioreagents | WRAB-9337 | 1:500 | AB_2335890 |
| Mouse anti-ABCA3 | Seven Hills Bioreagents | WMAB-17G524 | 1:500 | AB_577285 |
| Mouse anti-AGER | Abcam | ab54741 | 1:250 | AB_2242462 |
| Rabbit anti-PDPN | Santa Cruz Biotechnology | sc-134482 | 1:500 | AB_2162079 |
| Rabbit anti-HOPX | Santa Cruz Biotechnology | sc-30216 | 1:250 | AB_2120833 |
| Mouse anti-SFTPA | Leica | NCL-L-SPA | 1:200 | AB_564143 |
| Goat anti-TP63 | R&D Systems | BAF1916 | 1:500 | AB_2207173 |
| Mouse anti-HTII-280 | Terrace Biotechnology | TB-27AHT2-280 | 1:100 | AB_2832931 |
| Mouse anti-NAPSIN A | Leica | NCL-L-NAPSIN A | 1:500 | AB_10555426 |
| Mouse anti-MUC5AC | Abcam | ab79082 | 1:500 | AB_1603327 |
| Rabbit anti-KI67 | ThermoFisher | RM-9106-S1 | 1:500 | AB_2341197 |
| Rabbit anti-pSMAD1/5/8 | Millipore | AB3848 | 1:250 | AB_177439 |
| AffiniPure donkey anti-mouse IgG AlexaFluor 488 | Jackson ImmunoResearch | 715-545-150 | 1:500 | AB_2340846 |
| AffiniPure donkey anti-mouse IgG Cyanine3 | Jackson ImmunoResearch | 715-165-150 | 1:500 | AB_2340813 |
| AffiniPure<br>donkey anti-mouse IgG<br>AlexaFluor 647 | Jackson<br>ImmunoResearch | 715-605-150 | 1:500 | AB_2340862 |
| Secondary Antibodies |  |  |  |  |
| AffiniPure<br>donkey anti-<br>rabbit IgG<br>AlexaFluor 488 | Jackson<br>ImmunoResearch | 711-545-152 | 1:500 | AB_2313584 |
| AffiniPure<br>donkey anti-<br>rabbit IgG<br>Cyanine3 | Jackson<br>ImmunoResearch | 711-165-152 | 1:500 | AB_2307443 |
| AffiniPure<br>donkey anti-<br>rabbit IgG<br>AlexaFluor 647 | Jackson<br>ImmunoResearch | 711-605-152 | 1:500 | AB_2492288 |
| AffiniPure<br>donkey anti-<br>goat IgG<br>AlexaFluor 488 | Jackson<br>ImmunoResearch | 705-545-147 | 1:500 | AB_2336933 |
| AffiniPure<br>donkey anti-<br>goat IgG<br>Cyanine3 | Jackson<br>ImmunoResearch | 705-165-147 | 1:500 | AB_2307351 |
| AffiniPure<br>donkey anti-<br>goat IgG<br>AlexaFluor 647 | Jackson<br>ImmunoResearch | 705-605-147 | 1:500 | AB_2340437 |
| AffiniPure<br>donkey anti-mouse IgM | Jackson<br>ImmunoResearch | 715-545-140 | 1:500 | AB_2340845 |

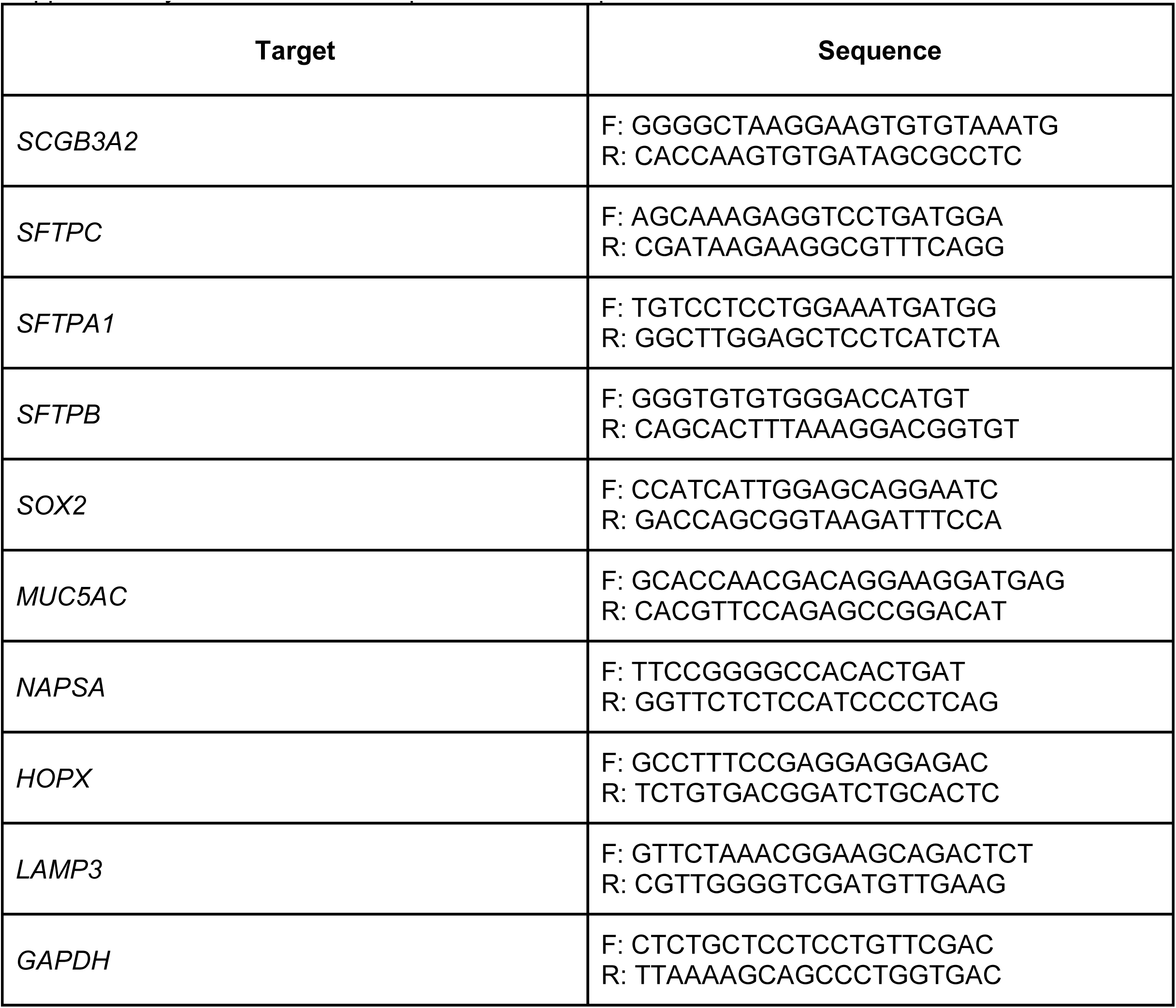
Supplementary Table 2. Primer sequences for RT-qPCR.

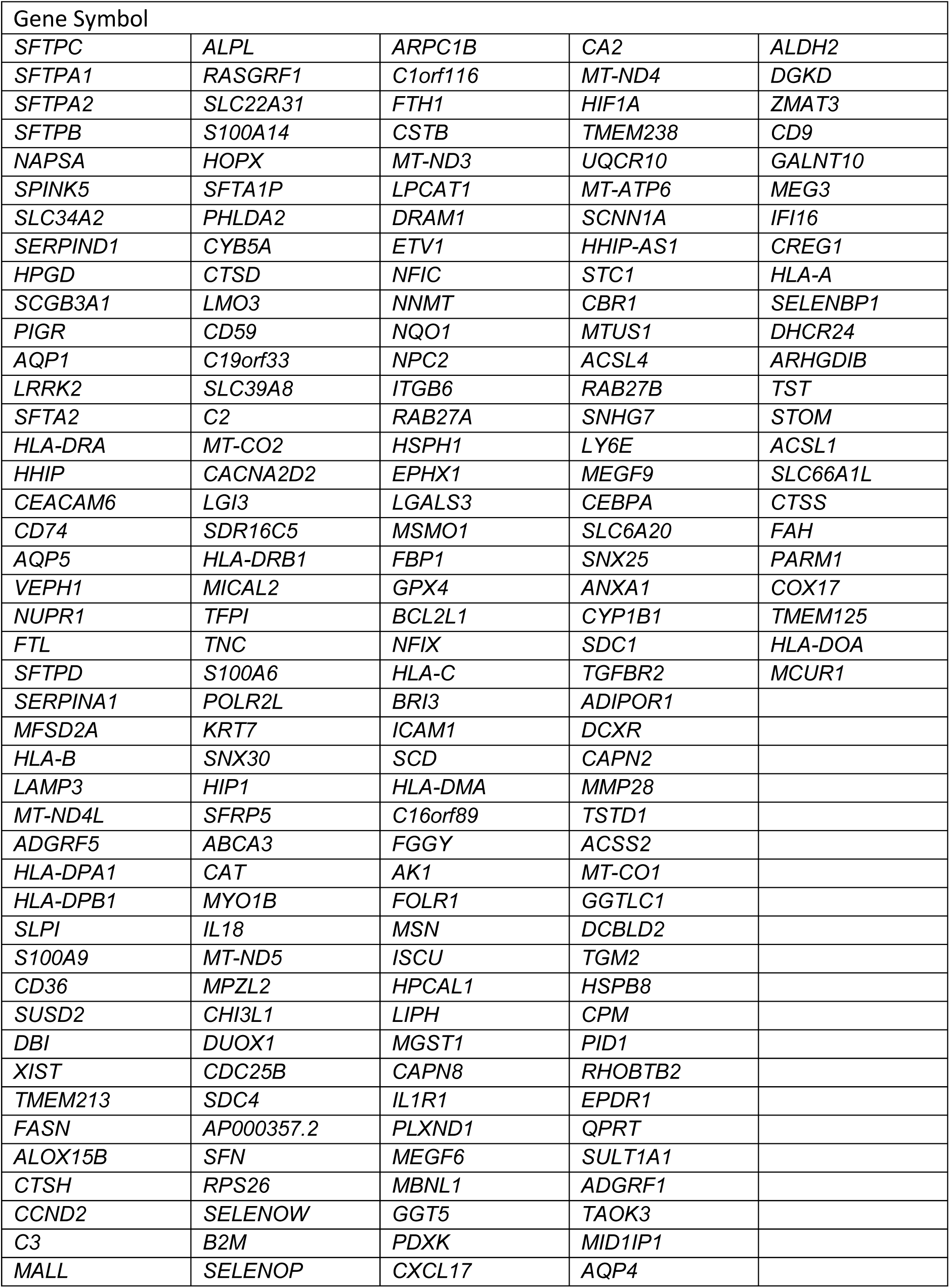
Supplementary Table 3: 199 genes enriched in primary alveolar type 2 organoids relative to bud tip progenitor organoids.

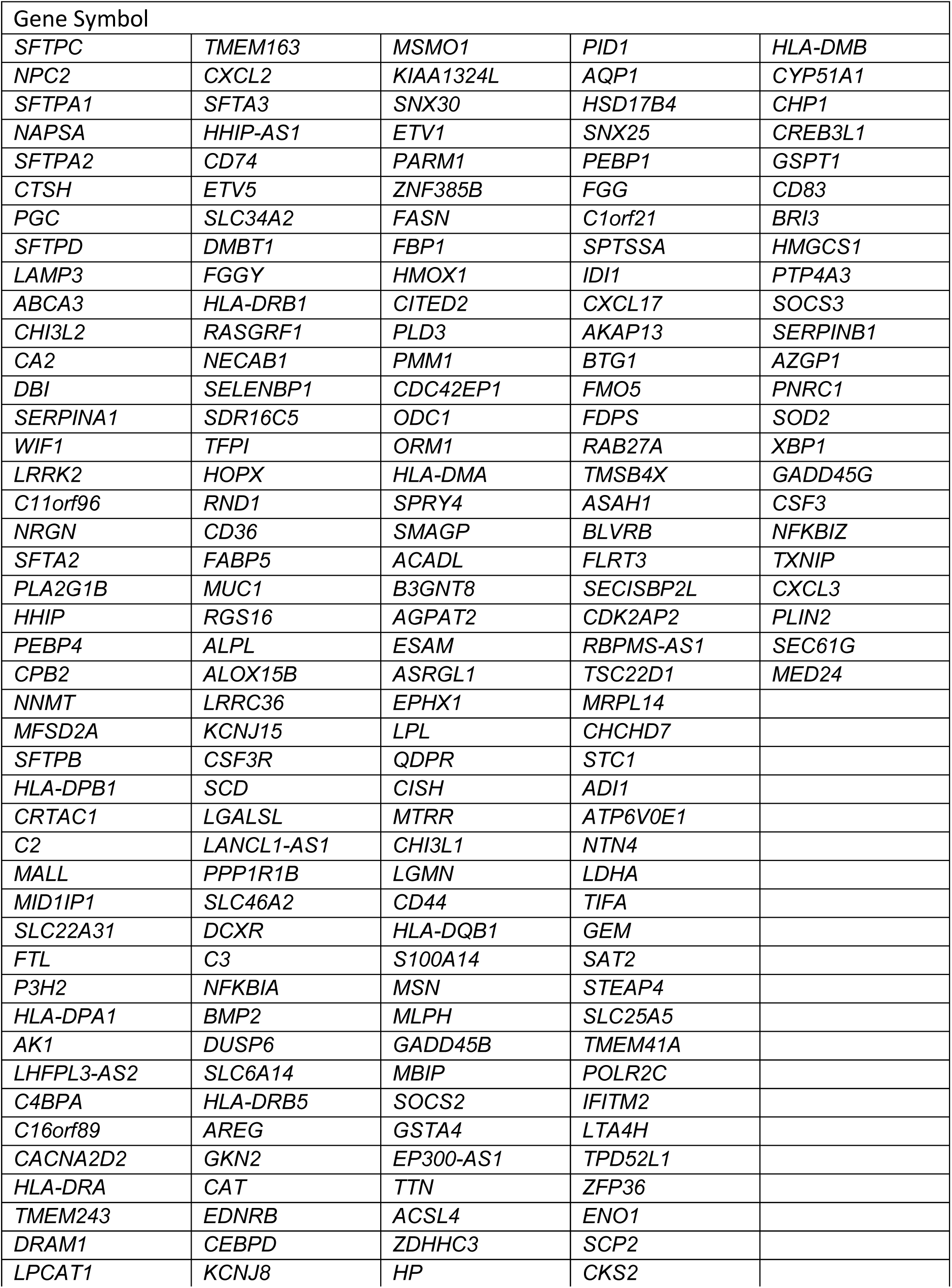
Supplementary Table 4: 199 genes enriched in primary alveolar type 2 cells relative to all other lung cell types. List from reference 27.

